# Stress-induced tyrosine phosphorylation of RtcB modulates IRE1 activity and signaling outputs

**DOI:** 10.1101/2020.03.02.972950

**Authors:** Alexandra Papaioannou, Federica Centonze, Alice Metais, Marion Maurel, Luc Negroni, Matías González-Quiroz, Sayyed Jalil Mahdizadeh, Gabriella Svensson, Ensieh Zare Golchesmeh, Alice Blondel, Albert C Koong, Claudio Hetz, Rémy Pedeux, Michel L. Tremblay, Leif A. Eriksson, Eric Chevet

**Author notes:** These authors contributed equally to the work (Alexandra Papaioannou, Federica Centonze). Correspondence to EC or LAE.

## Abstract

Endoplasmic Reticulum (ER) stress is a hallmark of various diseases, which is dealt with through the activation of an adaptive signaling pathway named the Unfolded Protein Response (UPR). This response is mediated by three ER-resident sensors and the most evolutionary conserved, IRE1α signals through its cytosolic kinase and endoribonuclease (RNase) activities. IRE1α RNase activity can either catalyze the initial step of XBP1 mRNA unconventional splicing or degrade a number of RNAs through Regulated IRE1- Dependent Decay (RIDD). The balance between these two activities plays an instrumental role in cells’ life and death decisions upon ER stress. Until now, the biochemical and biological outputs of IRE1α RNase activity have been well documented, however, the precise mechanisms controlling whether IRE1 signaling is adaptive or pro-death (terminal) remain unclear. This prompted us to further investigate those mechanisms and we hypothesized that XBP1 mRNA splicing and RIDD activity could be co-regulated by the IRE1α RNase regulatory network. We showed that a key nexus in this pathway is the tRNA ligase RtcB which, together with IRE1α, is responsible for XBP1 mRNA splicing. We demonstrated that RtcB is tyrosine phosphorylated by c-Abl and dephosphorylated by PTP1B. Moreover, we identified RtcB Y306 as a key residue which, when phosphorylated, perturbs RtcB interaction with IRE1α, thereby attenuating XBP1 mRNA splicing and favoring RIDD. Our results demonstrate that the IRE1α RNase regulatory network is dynamically fine-tuned by tyrosine kinases and phosphatases upon various stresses and that the nature of the stress determines cell adaptive or death outputs.

## Introduction

The imbalance between the cellular demand to fold secretory and transmembrane proteins and the Endoplasmic Reticulum (ER) capacity to achieve this function can result in the accumulation of improperly folded proteins in this compartment, a situation known as ER stress (Almanza *et al*, 2018). Activation of the ER stress sensors Inositol-Requiring Enzyme 1 alpha (referred to as IRE1 hereafter), Activating Transcription Factor 6 alpha (ATF6α) and Protein kinase RNA (PKR)-like ER kinase (PERK) aims to restore ER homeostasis and is known as the adaptive Unfolded Protein Response (UPR). However, when the stress cannot be resolved, the UPR triggers cell death (McGrath *et al*, 2021), which is referred to as terminal UPR. Thus far, the mechanisms controlling the switch between adaptive and terminal UPR remain incompletely characterized. Among the possible candidates, the IRE1 pathway being greatly conserved through evolution, plays crucial roles in both physiological and pathological ER stress. Similar to the other sensors, IRE1 is an ER transmembrane protein activated after dissociation from Binding immunoglobin Protein (BiP) and/or direct binding to improperly folded proteins (Karagöz *et al*, 2017).

IRE1 is characterized by the presence of both kinase and endoribonuclease domains in its cytosolic region. After its dimerization/oligomerization, IRE1 trans-autophosphorylates, allowing the recruitment of TRAF2 and subsequent activation of the JNK pathway (Urano *et al*, 2000). IRE1 dimerization and phosphorylation also yields a conformational change which activates its RNase domain and leads to unconventional splicing of the XBP1 mRNA and subsequent expression of a major UPR transcription factor XBP1s (Calfon *et al*, 2002; Lee *et al*, 2002). Importantly, the ligation following the IRE1-mediated cleavage of the 26 nucleotide intron in the XBP1 mRNA is catalyzed by the tRNA ligase RtcB (Lu *et al*, 2014; Jurkin *et al*, 2014; Kosmaczewski *et al*, 2014; Ray *et al*, 2014). IRE1 RNase domain also promotes the cleavage of mRNA (Hollien *et al*, 2009; Hollien & Weissman, 2006), rRNA (Iwawaki *et al*, 2001) and miRNA (Lerner *et al*, 2012; Upton *et al*, 2012) sequences, a process named Regulated IRE1-Dependent Decay (RIDD) of RNA (Hollien *et al*, 2009). Interestingly, the degree of oligomerization of IRE1α may define its RNase activity towards XBP1 mRNA splicing or RIDD, ultimately impacting on cell fate (Upton *et al*, 2012; Li *et al*, 2021; Le Thomas *et al*, 2021). Although recent and elegant direct approaches were used to measure the oligomeric changes that underpin IRE1 activation (Belyy *et al*, 2021), it remains unclear which is the order of oligomerization required for *XBP1* mRNA splicing vs. RIDD activity. However, there is a consensus on the cytoprotective effects of XBP1s and the cell death-inducing outputs of RIDD under acute ER stress (Tam *et al*, 2014; Upton *et al*, 2012; Han *et al*, 2009). RIDD was also described to maintain ER homeostasis under basal conditions (Maurel *et al*, 2014). In this model, while *XBP1* mRNA splicing is induced during the adaptive UPR (aUPR) and inactivated during the terminal UPR (tUPR), RIDD displays an incremental activation pattern reaching unspecific RNA degradation during tUPR. Despite our knowledge on IRE1 signaling biological outputs, little is known on the integration of these two RNase signals and how their balance impacts on the life and death decisions of the cell. The existence of a multiprotein complex recruited in IRE1 foci that dynamically changes composition during the course of ER stress, named “UPRosome”, has been suggested as a possible modulation mechanism (Hetz & Glimcher, 2009; Hetz & Papa, 2018). This is supported by the recent identification of IRE1 interactors (e.g. PP2A and TUBα1a) regulating XBP1 mRNA splicing differently from RIDD (Sepulveda *et al*, 2018).

Herein, we hypothesized that *XBP1* mRNA splicing and RIDD are part of an auto- regulatory IRE1 RNase network. RIDD targets could possibly impact on the splicing of *XBP1* mRNA, thus fine-tuning the response to ER stress and impacting on subsequent cell fate decisions. We identified *PTP1B* mRNA as a RIDD target affecting XBP1s activity by dephosphorylating RtcB, the tRNA ligase that performs *XBP1s* ligation. The tyrosine kinase c-ABL is also part of this network as it phosphorylates RtcB, thus revealing a post-translational regulatory mechanism of the IRE1 RNase signaling outputs. The study furthermore supports the true existence and dynamic nature of a UPRosome whose composition can determine the outcome of IRE1 signaling.

## Results

### PTP1B contributes to *XBP1* mRNA splicing and is a RIDD target

We first hypothesized that molecules involved in the IRE1-XBP1 pathway could at the same time be RIDD targets, and as such, could be part of a regulatory mechanism between both activities. To identify XBP1s regulators in the context of ER stress and possible RIDD substrates, we conducted two independent screens. First, a siRNA library against ER protein-coding RNAs was used to transfect HEK293T cells expressing a XBP1s-luciferase reporter (Spiotto *et al*, 2010) (**Fig. S1A**). The cells were then subjected to tunicamycin-induced ER stress, and luciferase activity was monitored to identify XBP1s positive and XBP1s negative regulators (**Fig. 1A**). We identified 23 positive and 32 negative regulators (out of a siRNA library targeting >300 genes) including candidates involved in metabolic processes (e.g. CH25H, DHCR7), post-translational modifications (e.g. POMT2, DMPK), ERAD pathway (e.g. SYVN1, UBC6) and cell growth (e.g. RRAS, CD74) (**Fig. 1B**). Second, to identify mRNA that could be cleaved by IRE1, we performed an *in vitro* IRE1 mRNA cleavage assay as described previously (Lhomond *et al*, 2018) (**Fig. S1B**) yielding a total of 1141 potential RIDD targets. XBP1s regulators that could also be subjected to RIDD-mediated degradation were determined by intersecting the hit lists from both screens, yielding 7 candidates, namely: ANXA6, ITPR3, PTPN1 (positive XBP1s regulators), RYR2, TMED10, KTN1 and ITPR2 (negative XBP1s regulators). The lists from this study were later combined with a list of XBP1s regulators coming from a genome-wide siRNA screen (Yang *et al*, 2018). Although we did not observe any common XBP1s modulators with our own study, likely explained by a sensitivity issue due to the targeted aspect of the library used in the present work, we note the existence of 33 more hits in the intersection between XBP1 splicing and RIDD. The functional association network of the reported candidates showed their involvement in UPR regulation, RNA processing and cellular homeostasis (**Fig. 1C-D, Fig. S1C**). Interestingly, among the 7 hits found was PTPN1, that encodes for the Protein-Tyrosine Phosphatase 1B (PTP1B), a protein previously reported by us to potentiate XBP1 splicing in response to ER stress (Gu *et al*, 2004).

**Figure 1.**
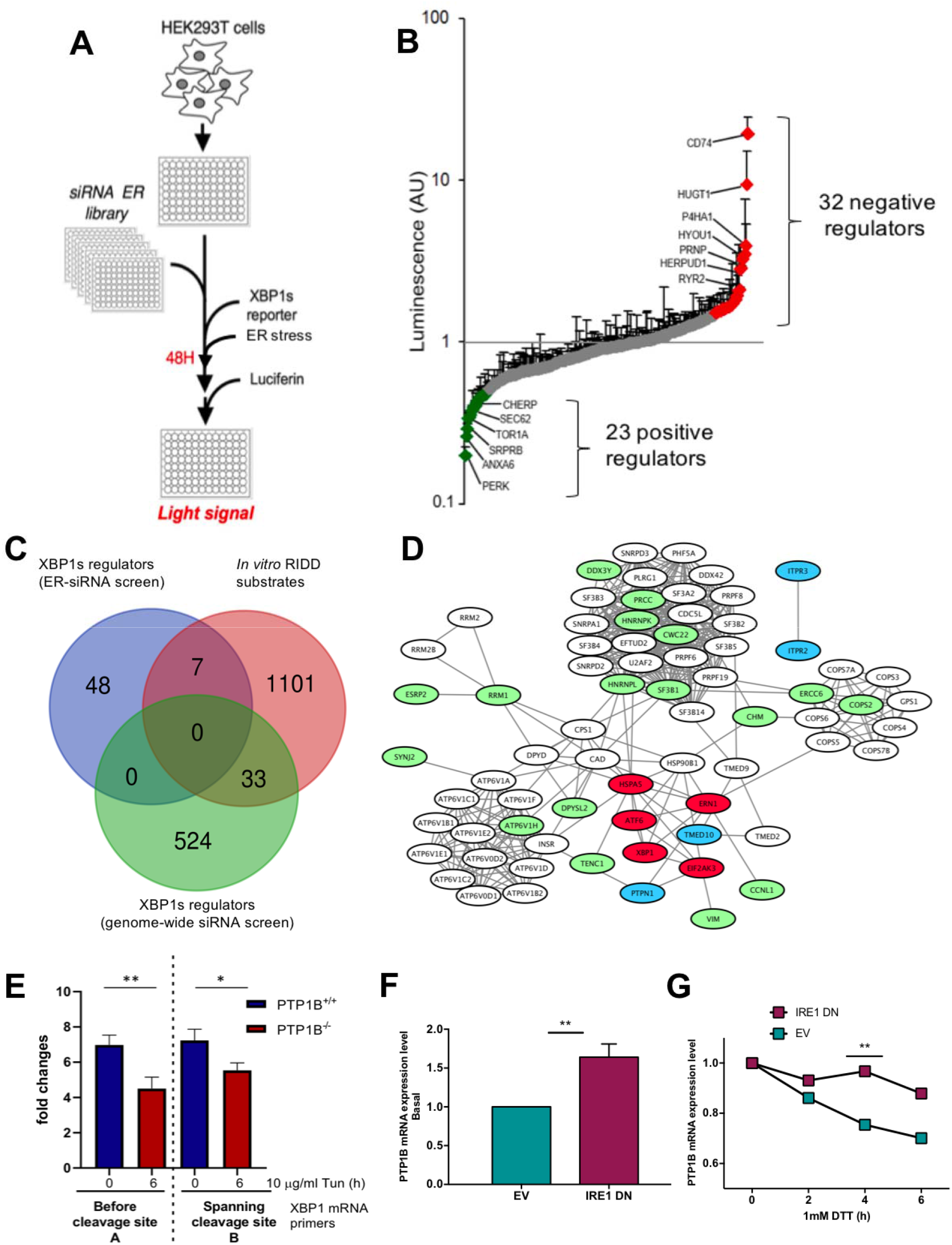
SiRNA screening and in vitro IRE1α cleavage assay identify PTP1B as both XBP1 splicing regulator and RIDD target. (**A**) siRNA-based screening assay: HEK293T cells were transfected with siRNA sequences against genes encoding ER proteins. They were subsequently transfected with an XBP1s-luciferase reporter (Fig. S1A) and after 48h and the induction of ER stress, the cells were tested for the intensity of light signal after the addition of luciferin. (**B**) Luminescence quantification of the screening assay described in (A). (**C**) Venn diagram of the gene list resulting from the assay described in (A-B), a list of possible RIDD substrates coming from an in vitro IRE1α cleavage assay (Fig. S1B) and a genome-wide siRNA-based screening assay (described in (Yang *et al*, 2018)). (**D**) A schematic network of the genes in the intersections of the lists as described in (C). (**E**) PTP1B^+/+^ (WT) and PTP1B^-/-^ (KO) mouse embryonic fibroblasts (MEFs) were treated for 0 and 6 hours with 10μg/ml TM. RNA was isolated and the resulting samples were analyzed with qPCR for spliced and total *XBP1* mRNA levels. The bar graph presents the tunicamycin-induced fold-change in *XBP1* mRNA splicing between PTP1B^+/+^ (WT, blue) and PTP1B^-/-^ (KO, red). ** indicates p = 0.0074; * indicates p = 0.0187. Total *XBP1* mRNA qPCR primers amplifying the region before the cleavage site or the region spanning the cleavage site were used (**Fig. S1C**). (**F-G**) PTP1B mRNA expression levels in U87 cells expressing or not a dominant negative form of IRE1α at 0h time-point (F) and during DTT treatment (1mM) with a 2-hour Actinomycin D (5 μg/ml) pre-treatment (G). EV: empty vector, IRE1 DN: dominant negative (cytosolic-deficient) form of IRE1α. Data information: Data values are the mean ± SEM of 4 independent experiments. Unpaired t test was applied for the statistical analyses.

To confirm our initial results we repeated these experiments by using RT-qPCR. PTP1B^-/-^ (KO) mouse embryonic fibroblasts (MEFs) exhibited a significantly lower capacity to yield *XBP1* mRNA splicing under tunicamycin-induced ER stress compared to their PTP1B^+/+^ (WT) counterparts (4-5 fold increase in PTP1B KO cells vs 7 fold increase in PTP1B WT cells; **Fig. 1E, Fig. S1D**). Regarding the UPR sensors ATF6 and PERK, the expression of both BiP and HERPUD was also attenuated in PTP1B^-/-^ cells while that of CHOP was increased, most likely as a compensatory mechanism (**Fig. S1E-G**). Using the same cellular settings, we also observed a basal activity of all three UPR sensors (**Fig. S1H**). We next sought to test whether PTP1B mRNA was indeed a RIDD substrate. To this end, we used U87 cells expressing a dominant negative form of IRE1α (IRE1 DN) or an empty vector (EV) (Drogat *et al*, 2007). We conducted an Actinomycin D chase experiment under basal or ER stress conditions to evaluate the post-transcriptional regulation of *PTP1B* mRNA expression. Under basal conditions, U87 DN cells showed higher expression levels of PTP1B compared to the control cells (**Fig. 1F**). Moreover, upon ER stress (DTT or Tunicamycin), *PTP1B* mRNA degradation was more efficient in control cells than in DN cells (**Fig. 1G, Fig. S1I**). This was a phenomenon also observable at the level PTP1B protein levels which significantly decreased upon ER stress compared to control cells (**Fig S1J**). Collectively, these results show that *PTP1B* mRNA is a genuine RIDD target and that PTP1B promotes IRE1-dependent *XBP1* mRNA splicing.

### RtcB is tyrosine-phosphorylated by c-ABL and dephosphorylated by PTP1B

Based on the experiments reported above, we concluded that PTP1B may represent a key regulator of IRE1 activity and investigated how this protein could mechanistically alter IRE1 signaling. PTP1B is a tyrosine phosphatase, and thus we first searched the PhosphoSitePlus database (Hornbeck *et al*, 2015) for the presence of phosphotyrosine (pY) residues in proteins described previously to regulate IRE1/XBP1s signaling. This revealed that while there was no reported pY residue for IRE1 (a result confirmed experimentally in our laboratory), six pY residues were reported for RtcB and found in phosphoproteomics studies (**Table S1**, (Hornbeck *et al*, 2015; Tsai *et al*, 2015; Bian *et al*, 2016)). Interestingly, multiple protein sequence alignment of the human RtcB and its orthologues in different species from the metazoan/animal kingdom revealed not only a high conservation of the protein across evolution, but also the conservation of specific tyrosine residues (**Fig. S2A**). To monitor RtcB tyrosine phosphorylation in cellular models, we transfected HEK293T cells with the pCMV3-mouse RTCB-Flag plasmid and treated them or not with the tyrosine-phosphatase inhibitor bpV(phen). Cell extracts were immunoprecipitated with anti-Flag antibodies, and the immune complexes were immunoblotted using anti-pY antibodies. BpV(phen) treatment augmented the pY signal in the whole cell lysates (**Fig. S2B**) and showed that endogenous RtcB was also subjected to tyrosine phosphorylation (**Fig. S2C**). We are aware that bp(V)phen treatment might create a bias in the analysis, but in this case systemic inhibition of protein tyrosine phosphatases was a pre-requisite for the visualization of pY-RtcB.

We then tested whether RtcB interacts with PTP1B. To this end, HEK293T cells were transfected with RtcB-Flag and either PTP1B-WT or PTP1B-C215S (trapping mutant; (Jia *et al*, 1995; Zhang *et al*, 2000)) expression plasmids. Cell extracts were immunoprecipitated using anti-PTP1B antibodies and the immune protein complexes were immunoblotted using anti-RtcB antibodies. RtcB-Flag exhibited a more stable association with the C215S PTP1B than what was observed for its wild-type form (**Fig. 2A**), suggesting that the pY residue(s) in RtcB might be trapped by the mutant PTP1B. To further document the functional relationship between PTP1B and RtcB tyrosine phosphorylation, mRtcB-Flag was transfected in PTP1B^-/-^ and PTP1B^+/+^ MEFs, and the RtcB-Flag pY status was evaluated as described above. This revealed that the mRtcB- Flag pY signal was increased in PTP1B^-/-^ MEFs when compared to PTP1B^+/+^ MEFs (**Fig. 2B**), which demonstrates that PTP1B is involved in the dephosphorylation of RtcB.

**Figure 2.**
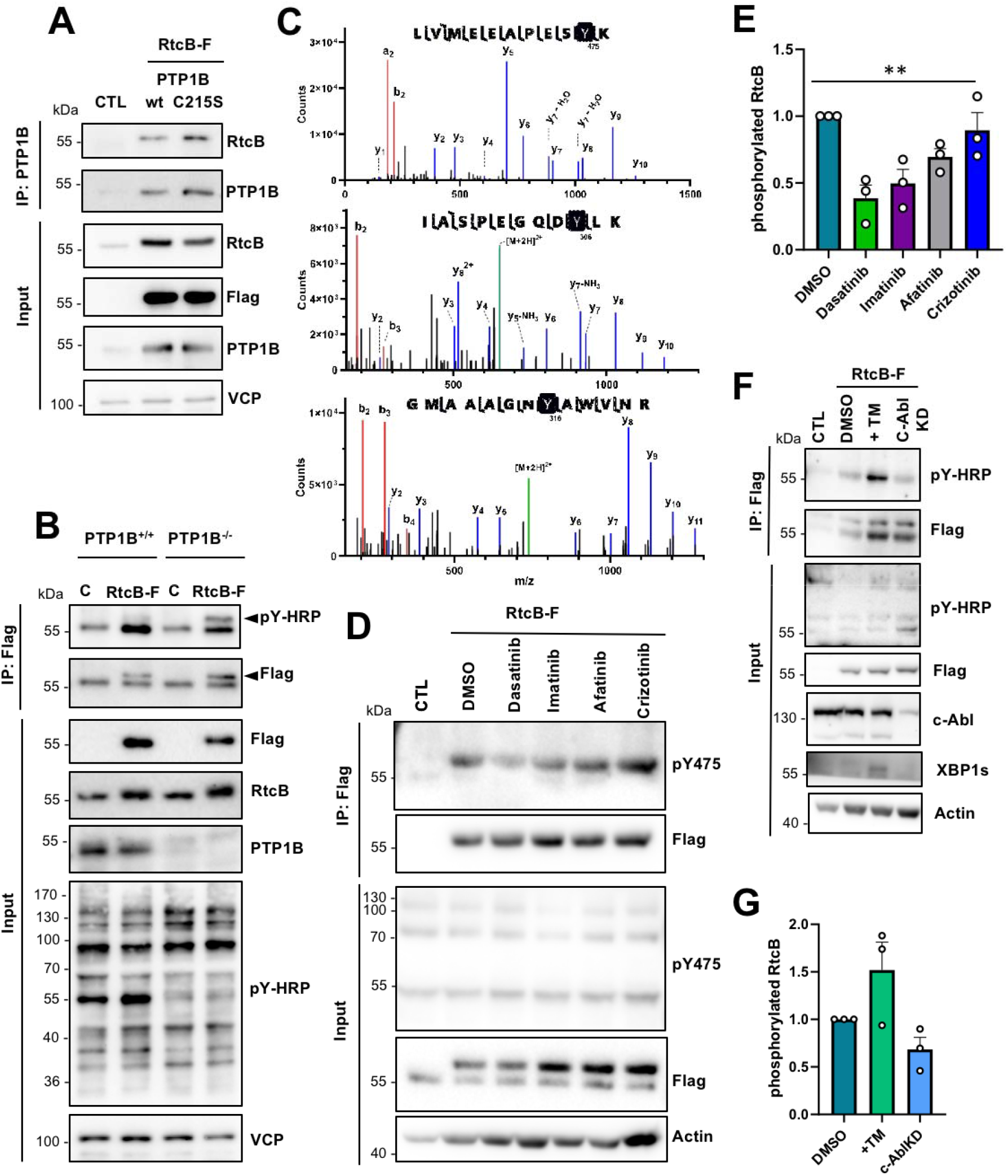
RtcB is a substrate of the tyrosine kinase c-Abl and the tyrosine phosphatase PTP1B. (**A**) HEK293T cells were left non-transfected (CTL) or transfected with 1μg of the wt RtcB-Flag and 1μg of either the wt PTP1B plasmid or the C215S mutant one. Immunoprecipitation was done in the cell lysates with the PTP1B antibody and the immunoprecipitates were immunoblotted for RtcB, while the membrane was re-probed with PTP1B. Inputs probed for RtcB, Flag, PTP1B and VCP are shown. (**B**) PTP1B^+/+^ or PTP1B^-/-^ MEFs were left untransfected (CTL) or transfected with 2μg wt RtcB-Flag and treated with 15μM bpV(phen) for 2 hours. Immunoprecipitation was done in the cell lysates using Flag Ab and the immunoprecipitates were immunoblotted for phosphotyrosine, while the membrane was re-probed with Flag. Input samples probed for Flag, RtcB, PTP1B, pY-HRP and VCP are shown. Black arrowheads indicate the RtcB-Flag protein, white arrowhead indicate an unspecific band at 55kDa. (**C**) Samples from a scaled-up in vitro kinase reaction containing recombinant human c-ABL and RtcB were analyzed using mass spectrometry. Fragment spectra corresponding to three different phospho-peptides containing tyrosine residues are depicted (y ions are shown in blue and b ions in red). (**D**) HEK293T cells transfected with the wt RtcB-Flag were treated 24 hours post-transfection with 10 μM of tyrosine kinase inhibitors afatinib, crizotininb, dasatinib and imatinib for 8 hours and 15 μM bpV(phen) for 2 hours. Immunoprecipitation was done in the cell lysates using Flag antibody and the immunoprecipitates were first immunoblotted for RtcB-pY475 and then re-probed with Flag Ab. Input samples probed for pY-HRP, Flag and VCP are shown. (**E**) The levels of phosphorylated RtcB (D) were normalized to RtcB protein levels (D). (**F**) HEK293T cells transfected with the Flag-RtcB-WT were treated 24 hours post- transfection with 1ug/ml TM for 6 hours or transfected with c-ABL siRNA for 2 days. Immunoprecipitation was done in the cell lysates using Flag antibody and the immunoprecipitates were first immunoblotted for pY and then re-probed with Flag antibodies. Input samples probed for pY-HRP, Flag, c-ABL, XBP1s and Actin are shown. (**G**) The levels of phosphorylated RtcB (F) were normalized to RtcB protein levels (F). Data information: The blots shown are representative of 3 or more independent experiments. Data shown in the graphs correspond to the mean ± SEM of n=3 independent experiments. One-way ANOVA was applied for the statistical analyses (**p < 0.01).

We next sought to identify a tyrosine kinase that could be responsible for RtcB phosphorylation. It was shown that the tyrosine kinase c-ABL acts as a scaffold for IRE1 by assisting its oligomerization and thus enhancing terminal UPR but is not directly involved in IRE1 phosphorylation (Morita *et al*, 2017). To document the possible phosphorylation of RtcB by c-ABL, we performed an *in vitro* kinase assay using recombinant human His-tagged c-ABL as kinase and recombinant human GST-RtcB as substrate. This experiment showed that c-ABL phosphorylated RtcB *in vitro* (**Fig. S2D**). We then mapped the tyrosine phosphorylation sites on RtcB using mass spectrometry and identified three tyrosine residues Y306, Y316 and Y475 that were subjected to phosphorylation (**Fig. 2C**). These tyrosines were also reported in independent proteomics screens to be phosphorylated (**Table S1**). Among them, the LVMEEAPESpY^475^K phosphopeptide was found not only in the TiO_2_ –purified but also in the crude sample indicating its probable greater abundance. In an attempt to use this phosphopeptide as a proxi of RtcB tyrosine phosphorylation, we developed an immunological tool for the selective detection of pY of RtcB at Y475, as described in **Fig. S3**. We observed the significantly reduced mRtcB-Flag tyrosine phosphorylation when cells were treated with the tyrosine kinase inhibitor dasatinib which targets c-ABL in a specific manner and which may also affect the activity of other tyrosine kinases such as SRC (**Fig. 2D**, **Fig. 2E**). Finally, we tested how RtcB tyrosine phosphorylation was affected upon ER stress and in c-ABL depleted cells (**Fig. 2F-G**). These experiments revealed that global tyrosine phosphorylation of RtcB-WT increased upon TM treatment and decreased in c-ABL knocked-down cells (**Fig. 2F-G**). This supports our finding that pY-mediated regulation of RtcB is an ER stress-related event.

### Molecular dynamics simulations of tyrosine phosphorylated RtcB and possible outcomes

We next evaluated the impact of tyrosine phosphorylation on RtcB structure using molecular dynamics (MD) simulations. The starting structure was a recent homology model of the human RtcB protein (Nandy *et al*, 2017). The systems that were prepared and subjected to MD simulations were distinguished by the presence (active) or absence (inactive) of bound GMP at the active site of RtcB, and by the different combinations of pY. After the simulations, certain conformational differences on the RtcB systems such as the movement of helices or specific amino acids were observed (**Fig. 3A, B, Fig. S4, 5**). By itself, the addition of a phosphoryl group to a tyrosine gives a negative charge to this modified amino acid, that in turn induces local changes to both the electrostatic surface and the conformation. Collectively, these alterations can have implications on protein structure, substrate specificity and interaction surface. To quantify these changes, we performed post-MD analyses including determination of the solvent accessibility surface area (SASA) and pKa values for the three tyrosine residues, as well as characterization of each tyrosine residue as “buried” within the protein structure or more solvent exposed (**Table S2, S3**). In addition, we quantified the dihedral angles for each of the three tyrosines in all the different RtcB systems as a measure of their rotation (**Fig. 3C**). We observed that when RtcB is fully phosphorylated, Y306 completely changes orientation to become more solvent exposed than buried in the body of the ligase (**Fig. 3D**, **Fig. 3A, B**). pY306 is characterized as a residue existing in both fully buried and solvent-exposed conformations (**Table S2**). Consistently, analysis of the final structures after the 200ns simulations revealed that, although the differences in the SASA values are not striking, pY306 has the highest value of SASA among the three phosphotyrosines (pY306/pY316/pY475), and a higher SASA value than Y306 in the non-pY system (**Table S3**). Next, we measured the docking score of GTP in the active site of RtcB in the differentially tyrosine phosphorylated systems. Phosphorylation of Y475 totally abrogated the binding of GTP in the active site (**Fig. 3E, F**). This is not surprising since Y475 is located at the active site of the protein and the bulky phosphoryl group creates a barrier that does not allow GTP to bind to the His 428 residue for the subsequent enzymatic catalysis.

**Figure 3.**
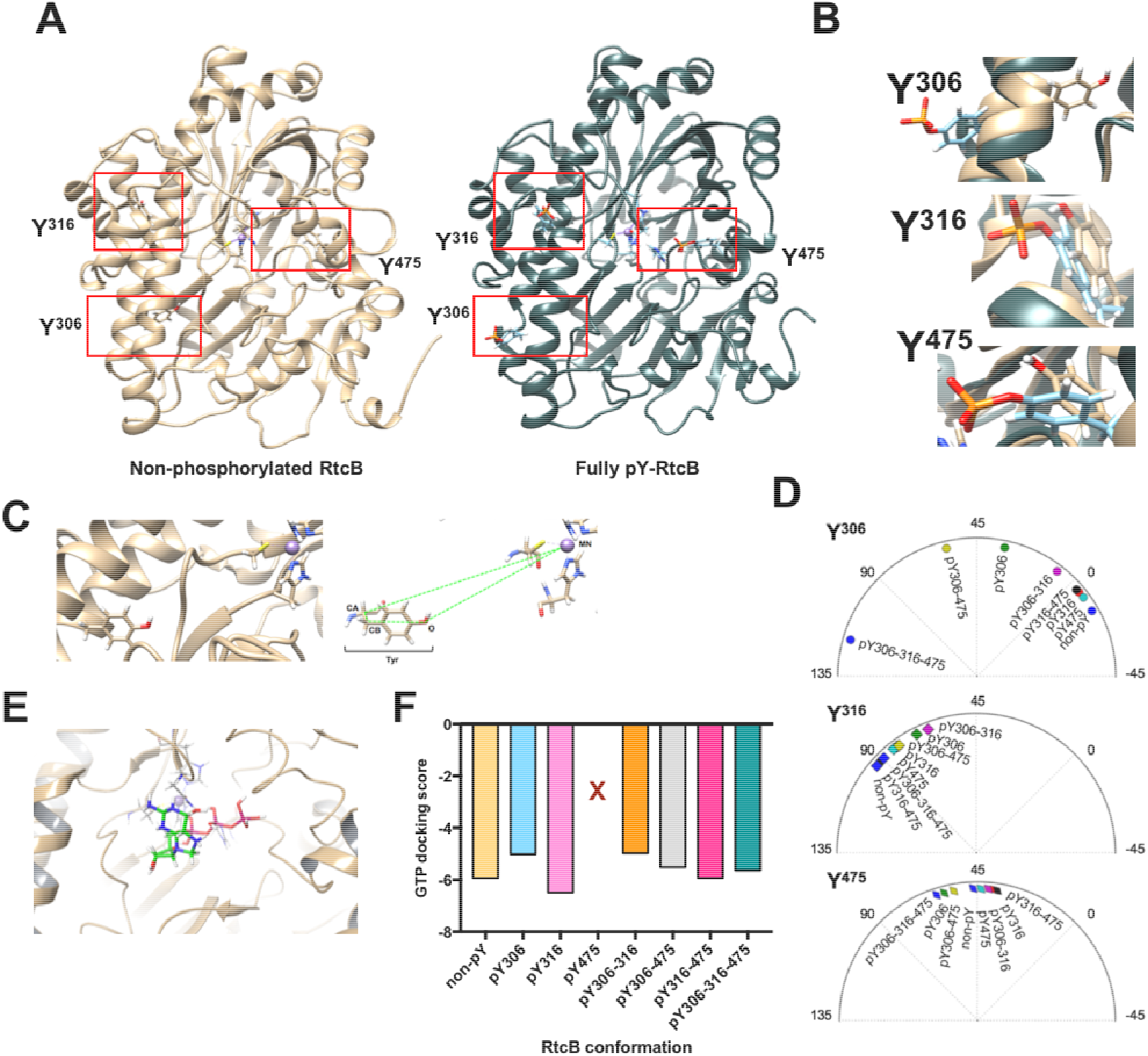
Molecular dynamics of RtcB tyrosine-phosphorylated systems and possible implications. (**A**) Ribbon-like structures of the RtcB protein resulting from the MD simulations either non-phosphorylated (left) or phosphorylated at Y306, Y316 and Y475 (right). (**B**) Zoom-in at Y306, Y316 and Y475 after superposition of the two RtcB models shown in (A). (**C**) Scheme explaining the calculation of a dihedral angle for a tyrosine relative to the Mn^2+^ in the active site of RtcB. The example shows Y306 in the unphosphorylated RtcB system. (**D**) The dihedral angles in degrees (°) of each of the three tyrosines 306, 316 and 475 relative to the metal ion in the protein active site in all RtcB systems after 200 ns MD simulation, depicted in polar charts. (**E**) Scheme showing the docked GTP ligand in the active site of the unphosphorylated RtcB protein. (**F**) Glide XP docking scores of GTP in the active site of the different RtcB systems. X represents the absence of GTP binding to the active site when Y475 is phosphorylated.

### Cellular impact of RtcB tyrosine phosphorylation

We then tested the effects of preventing phosphorylation of these tyrosine residues in a cellular environment. To this end, we generated RtcB-Flag Y-to-F mutants (**Fig. 4A**) using site-directed mutagenesis to keep the aromatic ring in a conformation that cannot be phosphorylated. Single (Y306F, Y316F, Y475F), double (Y306F/Y316F) and triple (Y306F/Y316F/Y475F) mutants were obtained. HEK293T cells were transfected with the WT and mutant plasmids, treated with bpV(phen) and lysates were then immunoprecipitated with anti-Flag antibodies. Anti-pY immunoblot revealed that while the Y306F and Y316F single mutants showed similar levels of tyrosine phosphorylation to WT, the Y475F mutant had a reduced degree of tyrosine phosphorylation indicating a potential preference for this residue by c-ABL (**Fig. 4B**). The double Y306F/Y316F mutant showed reduced tyrosine phosphorylation compared to the corresponding single mutants (**Fig. 4B**). The triple mutant still displayed tyrosine phosphorylation suggesting that other Tyr residues could be phosphorylated most likely by other tyrosine kinases. Quantitation of these Western blots indicated a decrease in tyrosine phosphorylation correlating with the number of tyrosine to phenylalanine conversion (**Fig. 4C**). Moreover, we tested how tyrosine phosphorylation of the different RtcB phospho-ablating mutants was affected upon TM-induced ER stress and in c-ABL-depleted cells as previously shown in **Fig. 2F-G**. Tyrosine phosphorylation of the RtcB mutants appeared, in general, to be less affected than that of RtcB-WT (**Fig. S6A**). This could be an indication that the phosphorylation events on single tyrosine residues studied herein are interconnected and might regulate each other. Of note, tyrosine phosphorylation monitored for RtcB-Y306F upon TM-induced ER stress exhibited an opposite behavior to that of RtcB-WT. Tyrosine phosphorylation of the RtcB-Y475F mutant appeared to be the least affected, supporting our observation that Y475 might be the most abundant pY in RtcB. Our results also showed that in c-ABL- depleted cells, RtcB-Y306F behaved similarly to RtcB-WT. In contrast, RtcB-Y316F showed an increased tyrosine phosphorylation which was the opposite of what was observed for both RtcB-WT and RtcB-Y306F. This observation might be indicative of the implication of other kinases phosphorylating these residues outside c-ABL. Furthermore, RtcB-Y475F appeared to be the least affected when c-ABL was depleted. Again as indicated above, this result supported our initial finding that Y475 might be the best c-ABL substrate (as reflected by its abundance) in RtcB (**Fig. S6A**). Collectively our results confirm that RtcB contains tyrosine residues substrates of c-ABL kinase activity. Moreover, they may indicate that the sequence of tyrosine phosphorylation on RtcB might impact on the relative phosphorylation of the different tyrosines. The additional pYs found in previous mass spectrometry-based analyses are shown in **Table S1**.

**Figure 4.**
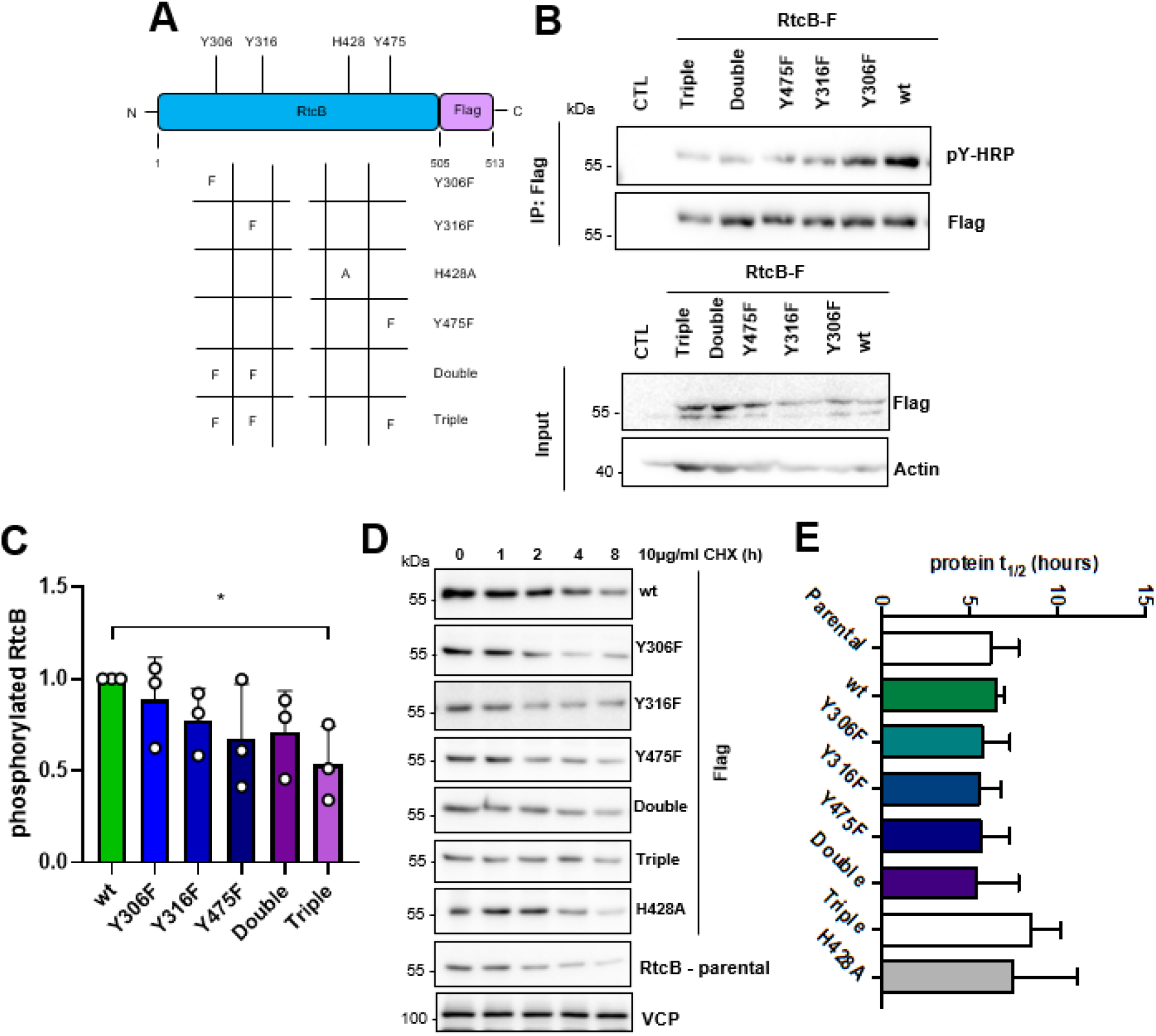
Generation of cellular models with altered RtcB tyrosine phosphorylation status. (**A**) Scheme showing the RtcB-Flag protein encoded by the pCMV3-mRtcB-Flag plasmid and the mutant forms that were created in this study. (**B**) HEK293T cells were left untransfected (CTL) or transfected with 2 μg of the different RtcB-Flag plasmids. 24 hours after transfection the cells were treated with 15 μM bpV(phen) for 2 hours. IP was performed in the cell lysates using Flag antibody and the immunoprecipitates were immunoblotted for phosphotyrosine. The membrane was re-probed with Flag. The corresponding input samples are also shown immunoblotted for Flag and Actin. (**C**) The levels of phosphorylated RtcB (B) were normalized to RtcB protein levels (B). (**D**) HeLa stable lines expressing wt or mutant RtcB-Flag were treated with 10μg/ml CHX for 0, 1, 2, 4 and 8 hours to block translation and monitor the current protein levels. Here, RtcB-Flag expression levels were monitored in the resulting protein lysates. The Parental HeLa cells were included to additionally monitor the levels of the endogenous RtcB. VCP was used as a loading control. (**E**) Half-life (t_1/2_) calculated for the different forms of the Flag-RtcB-WT protein or the endogenous RtcB in the case of the Parental HeLa cells. Data information: The blots are representative of 3 independent experiments. Data presented in the graphs correspond to the mean ± SEM of 3 independent experiments. One-way ANOVA was applied for the statistical analyses (*p < 0.02).

The interaction of each RtcB mutant with the substrate trapping mutant PTP1B- C215S was also tested as described earlier for RtcB-WT in **Fig. 2A**. The double (Y306F/Y316F) and mostly the triple (Y306F/Y316F/Y475F) mutants showed a statistically significant reduction of their physical association with the PTP1B trapping mutant compared to RtcB-WT (**Fig. S6B, C**). This could be indicative of a lack of preference for PTP1B to bind to one of the tyrosine sites but rather that the tyrosine phosphatase is able to bind and dephosphorylate all three sites Y306, Y316 and Y475. Following this observation, we generated HeLa cell lines stably expressing either the wt or the above-described mutant forms of RtcB-Flag (including also the catalytically inactive mutant H428A) (**Fig. 4A**). Cycloheximide chase-experiments were conducted on these cells to compare the protein stability of each RtcB-Flag forms. These analyses revealed that all RtcB-Flag proteins display a similar half-life (**Fig. 4D-E, S7A-C**) which was comparable to that of endogenous RtcB (**Fig. 4D-E, S7D**).

### Phosphorylation of RtcB at Y306 attenuates *XBP1* mRNA splicing and enhances RIDD activity

Since we confirmed that RtcB could be phosphorylated in cells, we next tested the impact of these phosphorylation sites on *XBP1* mRNA splicing. To this end, we treated HeLa stably expressing RtcB-Flag mutants cell lines with 1μg/ml Tun for 0, 2, 4, 8, 16 and 24 hours to monitor the *XBP1* mRNA splicing (**Fig. 5A, D**), and the results were normalized to the expression of RtcB-Flag (**Fig. 5C, F**), thereby indicating RtcB-Flag specific activity with respect to *XBP1* mRNA splicing (**Fig. 5B, E**). We observed that cells expressing the Y306F RtcB-Flag showed increased *XBP1* mRNA splicing activity (**Fig. 5A, B**). Furthermore, cells expressing the double mutant exhibited reduced XBP1 mRNA splicing compared to RtcB- WT-Flag but performed slightly better than cells expressing the catalytically inactive mutant H428A (**Fig. 5D, E**). The latter matched MD simulations results indicating that phosphorylation of RtcB on Y475 might abrogate the binding of GTP in the active site of the protein, thereby halting its catalytic activity. Interestingly, cells expressing the triple mutant showed a slightly lower XBP1 mRNA splicing than those expressing the H428A mutant thereby suggesting that the complete absence of c-ABL-mediated pY negatively affects RtcB in its contribution to *XBP1* mRNA splicing (**Fig. 5D, E**). Because of the observed effect of Y306F RtcB on *XBP1* mRNA splicing, we wondered whether RIDD activity would change as well. To address this, we performed an Actinomycin D chase experiment under ER stress (induced by either Tun or DTT) and monitored the expression *PER1* and *SCARA3* mRNAs, two known RIDD substrates whose expression levels decreased upon Tunicamycin-induced ER stress (**Fig. S7E**). Upon ER stress Y306F RtcB- Flag-expressing cells showed higher levels of *PER1* and *SCARA3* mRNA compared to control cells (**Fig. 5G**) corresponding to a lower RIDD activity. The same was observed even when the endogenous RtcB protein was knocked-down using a siRNA targeting the 3’-UTR of the *RTCB* mRNA (**Fig. 5G, H**). Despite the changes in *XBP1* mRNA splicing and RIDD activity, when the splicing of the three intron-containing tRNA molecules in human (corresponding to the Tyr, Arg and Ile tRNA) was compared between the Y306F and the RtcB-WT-Flag-expressing cells, there was no change in tRNA splicing activity (**Fig. 5I**). As a control to this experiment the levels of the intron-less tRNA molecules Pro and Val were also tested (**Fig. 5J**). These results indicated that phosphorylation of RtcB on Y306 represents a regulatory mechanism to control the balance between *XBP1* mRNA splicing and RIDD activity downstream of IRE1 RNase. The specificity of this regulation is consistent with the fact that tRNA splicing is not influenced by the RtcB Y306F mutation.

**Figure 5.**
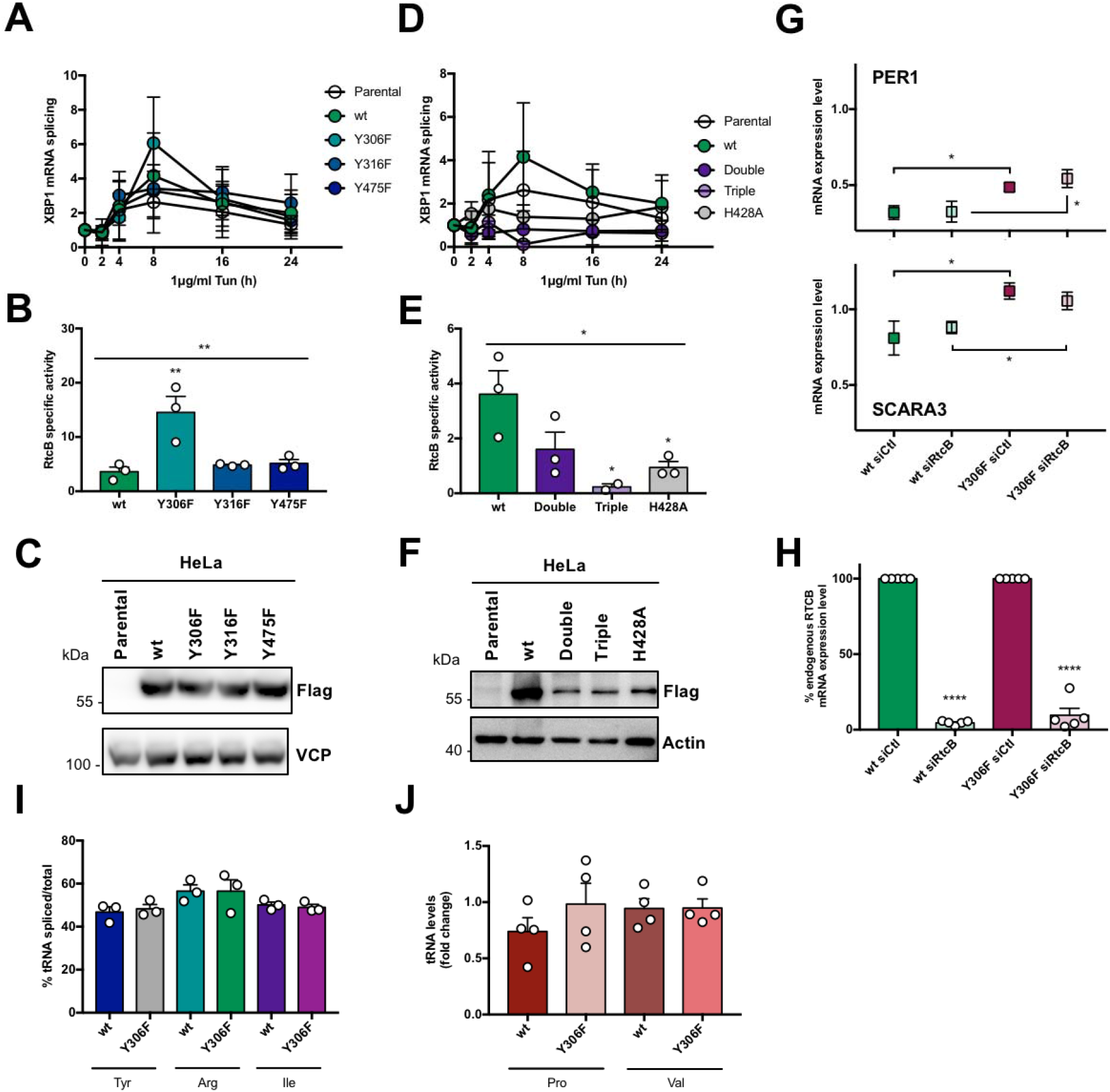
Phosphorylation of RtcB on Y306 negatively affects XBP1 mRNA splicing but not RIDD activity. **(A-F)** HeLa lines stably expressing wt or mutant RtcB-Flag were treated with 1μg/ml Tun for 0, 2, 4, 8, 16 and 24 hours. (**A, D**) cDNA coming from these was analyzed by qPCR for *XBP1s* and *XBP1* total mRNA levels with their ratio being the XBP1 mRNA splicing as depicted in the y axis of the graphs. The Parental HeLa cells are also included for comparison. (**B, E**) The peak of XBP1s activity at 8 hours (A, D) was then normalized to the protein levels (C, F) to obtain the RtcB specific activity. (**C, F**) Stable HeLa lines were lysed and analyzed for Flag-RtcB-WT expression levels by immunoblotting with anti-Flag. Actin and VCP were used as a loading controls. (**G**) HeLa lines stably expressing WT or Y306F Flag-RtcB were left untransfected or transfected with an siRNA targeting the 3’-UTR of the RTCB mRNA. 48 hours post-transfection the cells were pre-treated for 2 hours with 5μg/ml Actinomycin D and then treated either with 5μg/ml TM for 4 hours or 1mM DTT for 2 hours. The graph shows the mRNA levels of PER1 (upper part) and SCARA3 (lower part) after normalization to the untreated samples (0h time-point). (**H**) RT-qPCR analysis of the untreated samples from (G) for the endogenous RTCB mRNA using primers spanning its 3’-UTR. (**I**) cDNA from HeLa stable lines expressing wt or Y306F mutant RtcB-Flag was analyzed with qPCR for the splicing of the three intron-containing tRNA molecules in human Tyr, Arg and Ile. The graph shows the percentage of each spliced tRNA molecule to their total levels (unspliced + spliced). The presented values have been normalized to the levels of % tRNA splicing in Parental HeLa cells. (**J**) The levels of the intronless tRNA molecules Pro and Val were measured by RT- qPCR in cDNA samples from HeLa stable lines expressing wt or Y306F mutant RtcB-Flag. Values have been normalized to the levels measured in Parental HeLa cells. Data information: Data values for the graphs in (A-B), (D-E) and (I) are the mean ± SEM of 3 independent experiments. In (B) and (E) each mutant was compared to the wt using ordinary one-way ANOVA (*p < 0.05, **p < 0.01). Data values for the graphs in (G-H) and (J) are the mean ± SEM of, respectively, 5 and 4 independent biological experiments. Unpaired t-test was applied for the statistical analyses in those panels (*p < 0.05, ****p < 0.0001).

### Non-phosphorylable RtcB Y306 (Y306F) compensates defective *XBP1* mRNA splicing and confers resistance to ER stress-induced cell death

Based on the observations obtained from i) MD simulations that pY306 is almost totally exposed to the solvent and not buried in the protein in its fully-pY status, and ii) from the XBP1 splicing activity time-course of the respective HeLa line that Y306F RtcB-Flag contributes to more efficient *XBP1* mRNA splicing, we next tested if the phosphorylation of Y306 impacted on the interaction of RtcB with IRE1. This interaction was previously shown to ensure proper *XBP1* mRNA splicing (Lu *et al*, 2014). HEK293T cells were co-transfected with IRE1-WT and either WT or mutant forms of RtcB-Flag expression plasmids. Twenty-four hours post-transfection, cell lysates were immunoprecipitated with anti-Flag antibodies and immunoblotted with anti-IRE1 antibodies. We thus confirmed the interaction between IRE1 and RtcB (**Fig. 6A**) and found that the Y306F RtcB-Flag mutant exhibited a stabilized interaction with IRE1 compared to the WT form (**Fig. 6A**). The interaction of RtcB-WT and its pY306 variant with IRE1 and *XBP1* mRNA were then modeled (**Fig. 6B**). In the absence of XBP1, protein docking showed that RtcB-WT was able to bind essentially anywhere on the IRE1 tetramer surface with no specific referred site of interaction. However, upon *XBP1* mRNA binding, RtcB-WT docking unveiled two symmetrically positioned sites of interaction, one at each dimer, close to the corresponding *XBP1* mRNA splicing site (**Fig. 6B, top panels**). RtcB is oriented such that Y306 does not interact explicitly with IRE1/XBP1, whereas the catalytic area of RtcB is facing directly towards the cleavage site. Upon phosphorylation of Y306, RtcB instead binds in a position between the two cleavage loops of XBP1, at the IRE1 dimer-dimer interface (**Fig. 6B, lower panels**). The phosphate group of pY306 is pointing towards/interacting with *XBP1* mRNA, and the catalytic region of RtcB is facing directly towards the IRE1 tetramer with no direct interaction towards the *XBP1* mRNA. The free energies of binding as computed using MM-GBSA theory is similar for both WT and pY306 RtcB, albeit slightly stronger for the non-phosphorylated variant (−50.2, and −45.7 kcal mol^-1^, respectively). These docking calculations display a clear difference in the interaction of RtcB vs. pY306 RtcB, with the WT system being perfectly oriented so as to interact with the spliced ends of *XBP1* mRNA upon cleavage. Since our observations suggested that the RtcB Y306-dependent interaction could control the *XBP1* mRNA splicing activity of the IRE1/RtcB complex, we thus reconstituted *in vitro* the splicing reaction using *in vitro* transcribed *XBP1* mRNA, recombinant IRE1 cytosolic domain whose optimal concentration was empirically determined (**Fig. S7F**) and RtcB immunoprecipitated from cells. This reconstitution assay showed that as anticipated from our previous results, IRE1/RtcB Y306F spliced *XBP1* mRNA more efficiently than RtcB-WT (**Fig. 6C-E**). This result urged us to test whether the Y306F mutant would rescue the defective XBP1 splicing observed in PTP1B^-/-^ MEFs (**Fig. 1F, Fig. S8A, B**). PTP1B^-/-^ and PTP1B^+/+^ MEFs were either left untransfected or transfected with RtcB-WT-Flag or Y306F RtcB-Flag plasmids (**Fig. S8D**) and monitored for *XBP1* mRNA splicing upon 0, 2, 4, 8, 16 and 24 hours of 50ng/ml Tun treatment (**Fig. 6F**). Both WT and Y306F RtcB-Flag rescued XBP1 mRNA splicing in PTP1B^-/-^ MEFs. Importantly, quantification of these results revealed an better rescue of the Y306F RtcB- Flag consistent with the effect on the XBP1s activity observed in stable HeLa cell lines (**Fig. 6F, G**). The rescuing effect of Y306F RtcB-Flag was profound even under acute ER stress induced by 10μg/ml Tun for 6 hours (**Fig. S8C**). The importance of the phosphorylation event on the Y306 site was also reinforced by the docking of c-ABL on RtcB using MD simulation (**Fig. S8E**). The interacting residues on c-ABL (D325, D444) and RtcB (K279, R283, K357) created a surface of contact where Y306 of RtcB is well accessible to the active site of the kinase (**Fig. S8F**).

**Figure 6.**
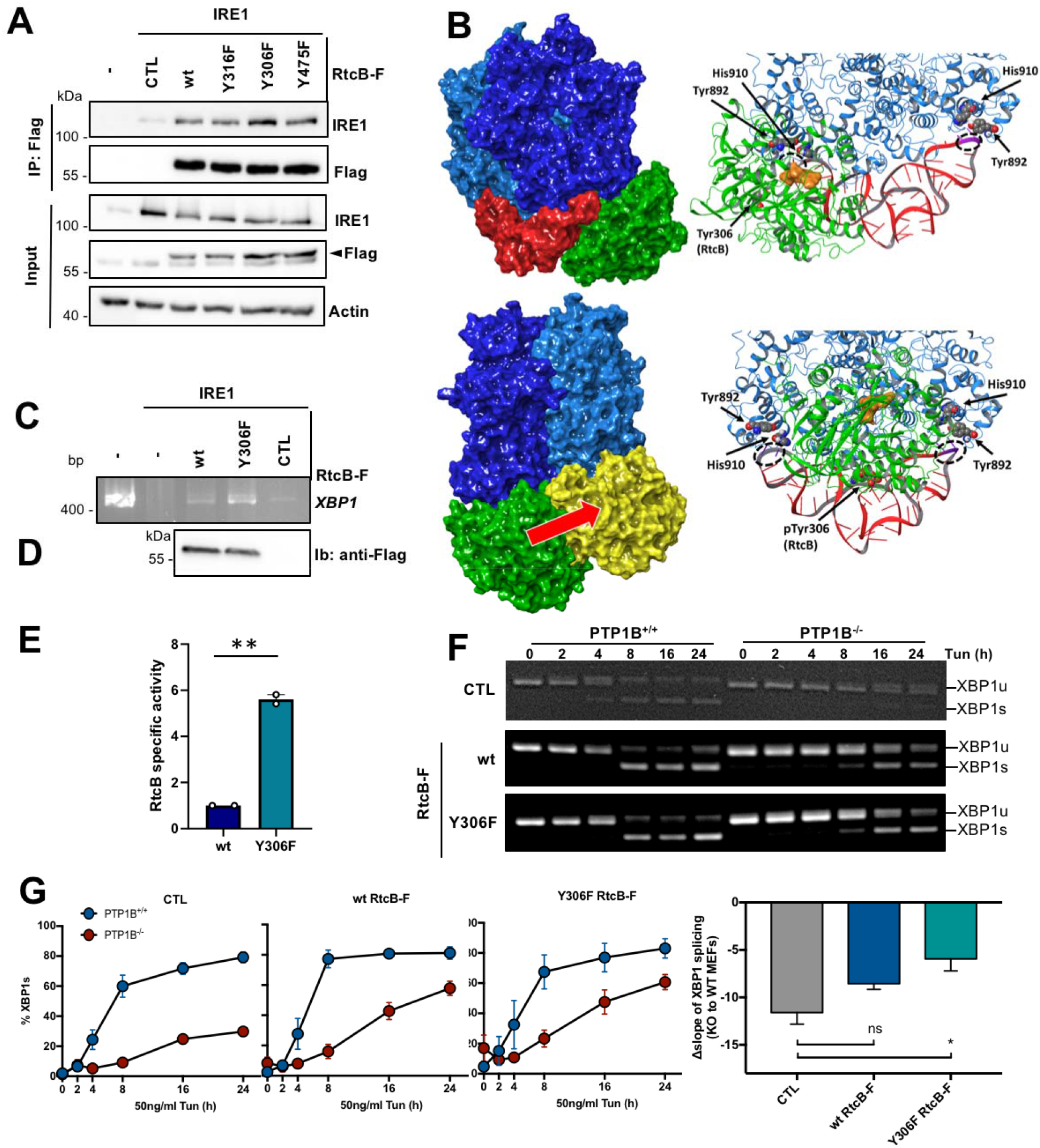
The Y306F RtcB mutant rescues a defective XBP1 mRNA splicing. (**A**) HEK cells were co-transfected with 1μg of wt IRE1α and 1μg of WT. or mutant Flag-RtcB plasmid. 24 hours later, cell lysis and anti-Flag immunoprecipitation was performed. The immunoprecipitates were blotted for IRE1α and the input samples for IRE1α, Flag and Actin. The arrowhead denotes the Flag-RtcB protein. (**B**) Docked complexes of RtcB WT and pY306 variants towards the IRE1 tetramer/XBP1 complex. (Top panels) RtcB WT (green) binds at the loop-binding RNase area of IRE1 (blue) and is perfectly oriented to initiate the ligation of XBP1 (red) upon its cleavage by IRE1. (Lower panels) The interaction area of the phosphorylated RtcB is shifted towards the middle of XBP1 at the IRE1 dimer-dimer interface region, and will not be able to initiate the ligation reactions of the spliced ends at the two XBP1 loops (pY306 of RtcB is represented in yellow). The red arrow indicates RtcB position shift on the IRE1 tetramer when Y306 is phosphorylated. (**C**) *In vitro* reconstitution of *XBP1* mRNA splicing using *in vitro* transcribed *XBP1* mRNA which was incubated with or without 250 ng of recombinant IRE1 cytosolic domain and with WT or Y306F RtcB-Flag immunoprecipitated from cells. The retrotranscribed XBP1 DNA from the assay was analyzed by PCR using primers recognizing the XBP1 mRNA. (**D**) Immunoprecipitated RtcB levels were detected by immunoprecipitating the cell lysates using anti-Flag antibodies conjugated beads and immunoblotting of the immunoprecipitates with anti-Flag antibodies. (**E**) XBP1s (C) was then normalized to RtcB protein levels (D) to obtain the RtcB specific activity. (**F**) PTP1B^+/+^ (WT) and PTP1B^-/-^ (KO) MEFs untransfected (CTL) or transfected with 2 μg of Flag-RtcB-WT or Flag-RtcB-Y306F plasmid were treated 24 hours post-transfection with 50ng/ml Tun for 0, 2, 4, 8, 16 and 24 hours. Their cDNA was analyzed by PCR using primers recognizing the *XBP1* mRNA. (**G**) Quantification of the gels in (F). The graph shows the comparison of XBP1 mRNA splicing in CTL cells and cells expressing either Flag-RtcB-WT or Flag-RtcB-Y306F. The composite comparison of the results obtained for ctl (circles) or rescues with either wt RtcB (triangles) or Y306F RtcB (squares) is also shown. (Bar graph) The slope of each curve was calculated and for each condition the slope of the WT cells was subtracted from the one of the KO cells, called as Δslope. The calculation of the Δslope were also corrected by the expression levels of Flag-RtcB-WT or Flag-RtcB-Y306F as determined by western blotting. Data information: Data values presented in (E) represent 2 independent experiments. Data values in (G) are the mean ± SEM of 3 independent experiments. Unpaired t-test was applied for the statistical analyses (ns: non-significant, *p < 0.05, **p < 0.01).

On the grounds of the pro-survival character of *XBP1* mRNA splicing, we then wondered whether phosphorylation of Y306 in RtcB could have an effect on the cellular survival/death. When cells stably over-expressing the WT or the Y306F form of RtcB were treated for 24 hours with increasing concentrations of DTT to induce ER stress, the survival advantage of the latter cells became apparent (**Fig. 7A**). Y306F RtcB-Flag expressing cells presented even less necrosis when endogenous RtcB was knocked-down by using an siRNA targeting the 3’-UTR of the *RtcB* mRNA. This meant that the observed effect was solely due to the exogenously expressed Y306F RtcB mutant (**Fig. 7A**). Under these specific stress conditions, in contrast to necrosis/late apoptosis, early cell apoptosis was not evident. Efficient silencing of endogenous RtcB was confirmed by both qPCR using primers spanning the 3’-UTR of the *RtcB* mRNA (**Fig. S8G**) and western blot analysis (**Fig. S8H**). These results indicate that RtcB Y306 is a key residue in the IRE1- XBP1/RIDD signaling since it is phosphorylated by c-ABL and dephosphorylated by PTP1B, and it regulates the interaction between RtcB and IRE1. Further to this, we realize that phosphorylation of RtcB on Y306 is able to direct a decision towards cell death in the presence of ER stress downstream of the IRE1 RNase signaling outputs. To further document the role of the pY-dependent interaction between RtcB and IRE1 in the control of cell death, we relied on the recent discovery that DNA-damaging agents signaled through RIDD (Dufey *et al*, 2020) and hypothesized that forcing the splicing of *XBP1* mRNA through the expression of Y306F RtcB should sensitize the cells to cell death. To test this, cells expressing wt or Y306F RtcB were treated with Doxorubicin or Etoposide and in both cases Y306F RtcB expressing cells were more prone to cell death than cells expressing wt RtcB (**Fig. 7B, S9 and S10**), thereby indicating that the balance between XBP1 mRNA splicing and RIDD i) is tightly controlled by RtcB, ii) plays a key role in stress-dependent life and death decisions and iii) determines the nature of the biological output depending on the stress cells are exposed to.

**Figure 7.**
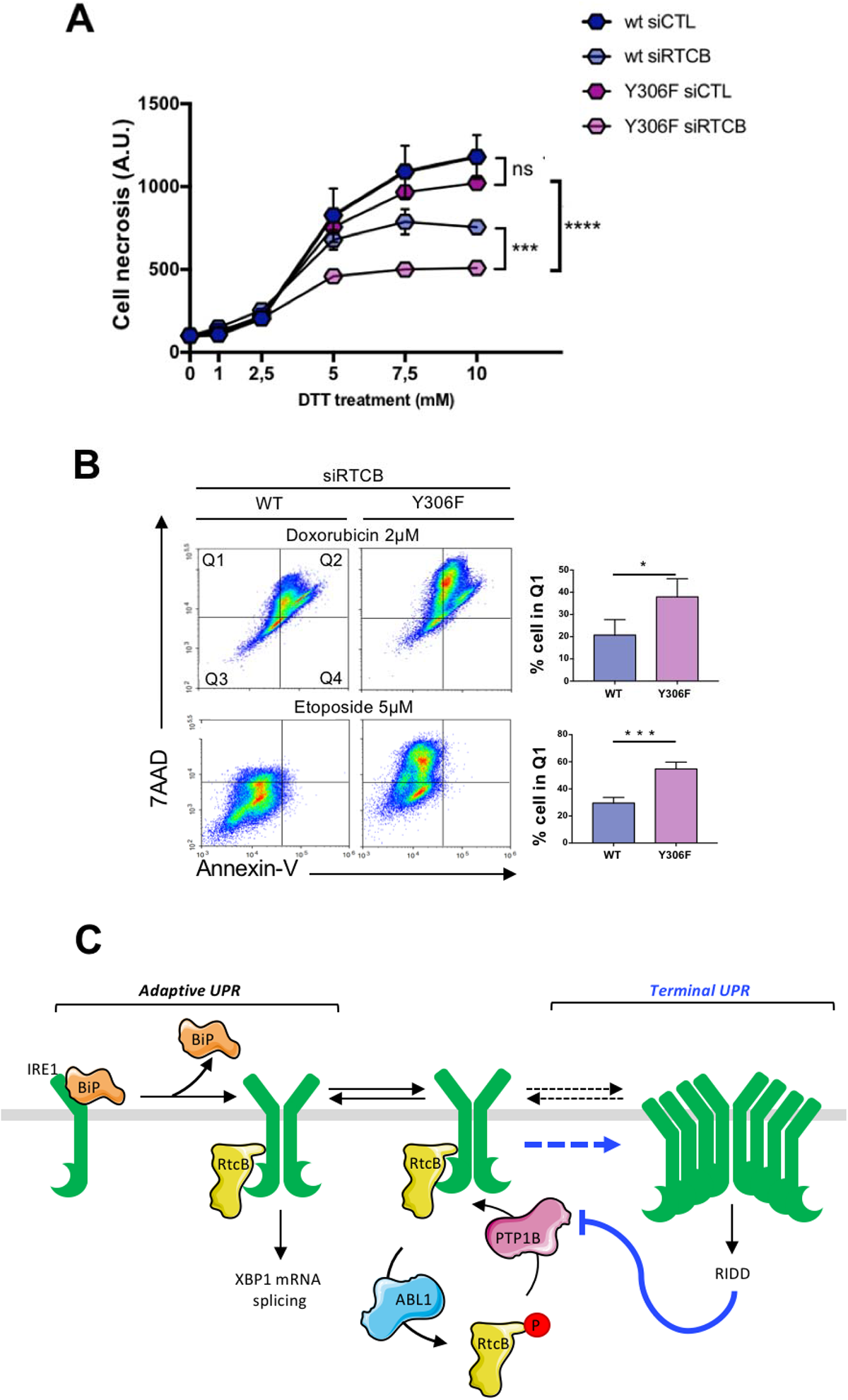
RtcB Y306 phosphorylation is a key event in modulating stress-induced cell life and death decisions. (**A**) HeLa lines stably expressing wt or Y306F RtcB-Flag were transfected or not with siRNA sequences against the endogenous RTCB mRNA. 48 hours post-transfection they were treated with 0, 1, 2.5, 5, 7.5 and 10 mM DTT for 24 hours. The resulting samples were then analyzed by FACS for cell necrosis and apoptosis through 7-AAD and Annexin V staining, respectively. Data values are the mean ± SEM of n=4 independent experiments. Two-way ANOVA and Tukey’s multiple comparisons test was applied for the statistical analyses (ns: non-significant, ***p < 0.001, ****p < 0.0001). (**B**) HeLa lines stably expressing Flag-RtcB-WT or Flag-RtcB-Y306F were left untransfected or transfected with siRNA sequences against the endogenous RTCB mRNA. 48 hours post-transfection they were treated with 0, 0.5, 1 and 2 μM Doxorubicin, or 0, 5, 10 and 20 μM Etoposide for 24 hours. The resulting samples were then analyzed by FACS for cell necrosis and apoptosis through 7-AAD and Annexin V staining, respectively. Data from 2 μM Doxorubicin and 5 μM Etoposide treatments are shown. Data values are the mean ± SEM of n = 4 independent experiments in two-way ANOVA and post-Tukey multiple comparisons test (*p < 0.05, ***p < 0.001). (**C**) Model representation of IRE1 activation and signaling towards XBP1s or RIDD. A decrease in PTP1B expression driven by RIDD is thought to reduce the formation of the IRE1/RtcB complex thereby pushing towards unleashed RIDD and terminal UPR. Dashed lines represent an higher order oligomerization of IRE1 which might result in terminal UPR.

## Discussion

The objective of this study was to better characterize how *XBP1* mRNA splicing and RIDD activities can crosstalk and how their signals are integrated. We showed that RtcB, the tRNA ligase responsible for *XBP1* mRNA splicing, is at the center of a kinase/phosphatase network and that the tyrosine phosphorylation-dependent regulation of RtcB interaction with IRE1 might represent a mechanism to shift IRE1 activity from *XBP1* mRNA splicing to RIDD. We have identified c-ABL as a tyrosine kinase causing RtcB phosphorylation on the three sites: Y306, Y316 and Y475, and PTP1B as a tyrosine phosphatase de- phosphorylating RtcB. Interestingly, the phosphorylation of Y475 on RtcB, a residue located in the active site, attenuated *XBP1* mRNA splicing. Indeed, the phosphorylation of Y475 prevents GTP binding to His 428, thus halting the process of ligation. More surprisingly, phosphorylation of Y306 weakened the interaction between RtcB and IRE1 thus reducing *XBP1* mRNA splicing activity. This could in turn favor the formation of IRE1 oligomers, leading to the observed increase in RIDD activity. In the same line of evidence, expression of RtcB-Y306F compensated defective *XBP1* mRNA splicing observed in PTP1B^-/-^ cells thereby indicating that phosphorylation on this specific residue impacts negatively on the *XBP1* mRNA splicing activity and constitutes a PTP1B-dependent mechanism in the UPR. Moreover, the fact that RtcB-Y306F conferred resistance to ER stress-induced cell death poses Y306 phosphorylation (preceding or following the phosphorylation on Y316 and Y475) as a key cellular event allowing life/death decisions (**Figure 7C**).

Our data confirm our initial hypothesis by demonstrating the existence of an auto- regulatory network emanating from the IRE1 RNase activity. In this regulatory network PTP1B was identified as a key actor by dephosphorylating RtcB, thereby facilitating the activation of the IRE1/XBP1s axis. *PTP1B* mRNA is a RIDD substrate such that when RIDD becomes dominant, PTP1B expression is prevented. Ultimately, one could propose a new model in which PTP1B would keep RtcB in a non-phosphorylated state thereby allowing the IRE1/RtcB complex to efficiently splice *XBP1* mRNA, hence exerting adaptive UPR. However, when the stress cannot be resolved, PTP1B expression would decrease through a RIDD-dependent mechanism. In parallel c-ABL is recruited to the ER membrane (Morita *et al*, 2017) where it phosphorylates RtcB on Y306, Y316 and Y475, thus further inhibiting IRE1/RtcB interaction initially lowered by the reduction of PTP1B expression, and in the meantime, enhancing RIDD by prompting IRE1 oligomerization, hence resulting in terminal UPR. Moreover, the relationship between IRE1 activity triggered by ER stress and IRE1 oligomeric state that was recently reported (Belyy *et al*, 2021) is in accordance with our model in which XBP1 mRNA splicing might be catalyzed by tetrameric oligomers. An integrated model could propose that ER stress-driven tyrosine phosphorylation of RtcB might lead to its dissociation from the IRE1 splicing complex which in turn would allow the formation of active IRE1 oligomers thus activating RIDD.

In line with these observations, it was demonstrated in a recent report that c-ABL can dissociate from 14-3-3 proteins in the cytosol upon ER stress, and be recruited to IRE1 foci at the ER membrane augmenting terminal UPR (Morita *et al*, 2017). Herein, we show that c-ABL can act not only as a scaffold for IRE1 but also through its catalytic activity with the phosphorylation of RtcB on tyrosine residues that in turn causes the reduction of its *XBP1* mRNA splicing activity. From our results and other studies, we can also speculate that apart from c-ABL, other tyrosine kinases might be involved in the regulation of the UPR. For instance, the SRC tyrosine kinase has recently been reported to be activated by and bind to IRE1 under ER stress leading to ER chaperone-relocalization to the cell surface (Tsai *et al*, 2018). As such, the scaffolding role of IRE1 could represent a means to integrate multiple signaling pathways thus far unrelated to IRE1 signaling, and link them to a clear function in cell homeostasis.

In the present study, we identified RtcB as a new target of the tyrosine phosphatase PTP1B. Interestingly, besides our initial report that PTP1B is involved in the control of IRE1 activity (Gu *et al*, 2004), PTP1B was subsequently found to also target the PERK arm of the UPR by negatively regulating its activation through direct dephosphorylation (Krishnan *et al*, 2011). As such, one might consider tyrosine kinase/tyrosine phosphatase signaling as a new key element in the regulation of the UPR to control life and death decisions. Our findings indicate that the exchange of a phosphatase for a kinase and vice versa is crucial for the *XBP1* mRNA splicing catalyzed by IRE1 and RtcB. In the molecular complex of IRE1 and RtcB, an additional member could potentially be the archease protein reported as a stimulatory co-factor of RtcB not only for tRNA but also for *XBP1s* mRNA ligation (Jurkin *et al*, 2014; Popow *et al*, 2014). Our results demonstrate that the dynamic alteration of the constituents of a multi-protein complex at the ER membrane can drive its collective activities, which further supports the notion of a UPRosome in the control of cellular life and death decisions (Hetz & Glimcher, 2009; Hetz & Papa, 2018).

## Materials and Methods

### Materials

Tunicamycin (Tun), dithiothreitol (DTT), Actinomycin D, cycloheximide (CHX) and sodium orthovanadate (Na_3_VO_4_) were obtained from Sigma-Aldrich. The phosphatase inhibitor bpV(phen) and the tysosine kinase inhibitors afatinib, crizotinib and sorafenib were from Santa Cruz Biotechnologies. The tyrosine kinase inhibitors dasatinib and imatinib (Imatinib Mesylate - STI571) were from Selleckchem. Protease and phosphatase inhibitor cocktail tablets were from Roche through Sigma-Aldrich. Pierce ECL Western Blotting Substrate was from Thermo-Fisher Scientific. The pCMV3-mRTCB-Flag plasmid encoding the wt form of mouse RtcB was purchased from Sino Biological. The various mutant forms of this plasmid were made with the QuikChange II XL Site-Directed Mutagenesis Kit from Agilent using the appropriate primers each time (**Table S4**). The pcDNA3.1/Zeo-PTP1B wt and pcDNA3.1/Zeo-PTP1B C215S plasmids were used as previously described (Blanchetot *et al*, 2005). The pCDH-CMV-IRE1α-EF1-Puro-copGFP was cloned in the lab as previously described (Lhomond *et al*, 2018).

### Cell culture and transfection

HEK293T, HeLa cells, U87 cells bearing a dominant negative form of IRE1α (IRE1 DN) or an empty vector (EV) (Drogat *et al*, 2007) and mouse embryonic fibroblasts (MEFs) wt or KO for PTP1B (obtained in (Gu *et al*, 2004)) were cultured in Dulbecco’s modified Eagle’s medium (DMEM) supplemented with 10% FBS at 37 °C in a 5% CO2 incubator. For the generation of HeLa cell lines stably expressing each one of the RtcB-Flag proteins, cells were first transfected using Lipofectamine 2000 (Thermo-Fisher Scientific) with 1μg of the corresponding plasmid. After 24 hours, the medium of the cells was changed to medium containing 600 μg/ml Hygromycin B (Thermo-Fisher Scientific) to start the selection process. The Hygromycin B-containing medium was replaced every three days for a total period of 21 days, when polyclonal populations of HeLa cells stably expressing each of the RtcB-Flag proteins were obtained. The stable cell lines were maintained in DMEM with 10% FBS containing 120 μg/ml Hygromycin B. Transient transfections were achieved using either polyethylenimine (PEI) for the HEK293T cells or Lipofectamine 2000 for the HeLa cells and the MEFs together with the desired plasmid. For preparation of the PEI solution, branched polyethylenimine (average M_w_∼25,000) powder was purchased from Sigma-Aldrich.

### siRNA-based screening assay

The library of approximately 300 small-interfering RNAs (siRNA) against genes encoding ER proteins was used in previous studies (Higa *et al*, 2014). Each siRNA (25 nM) was transfected into HEK293T cells by transfection with the Lipofectamine RNAiMAX reagent (Thermo-Fisher Scientific). The cells were then transfected with an XBP1s-luciferase reporter in which the firefly luciferase gene is fused to the XBP1 gene (Spiotto *et al*, 2010): under basal conditions where XBP1 mRNA remains unspliced, a stop codon at the gene fusion is in-frame thus not allowing the luciferase to be expressed. In contrast, under ER stress conditions the splicing of XBP1 mRNA shifts the ORF (open reading frame) resulting in the loss of the stop codon downstream of XBP1 and the subsequent expression of luciferase. ER stress was after applied to the cells and sequentially the substrate of the luciferase enzyme, luciferin. The cells were then tested for the emission of light that after appropriate quantification and a value threshold set-up led to the distinction of two major groups of genes: XBP1s positive and XBP1s negative regulators. High values of luminescence (>1 AU) signify that efficient XBP1 splicing took place so the luciferase expression was allowed, thus marking the silenced genes as negative XBP1s regulators. The opposite is true for values <1 AU that place the silenced genes in the category of positive regulators of XBP1 mRNA splicing.

### in vitro IRE1 mRNA cleavage assay

As described previously (Lhomond *et al*, 2018), in this assay total RNA was extracted from U87 cells, refolded and incubated or not with recombinant IRE1α protein under appropriate physicochemical conditions. The resultant RNA sequences from the cleaved and uncleaved conditions underwent polyA mRNA isolation, the final different groups (non-polyA: cleaved or uncleaved, and polyA: cleaved or uncleaved) were reverse transcribed and the respective cDNA material was hybridized on a human transcriptome array whose analysis revealed potential RIDD substrates.

### Immunoblot and Immunoprecipitation

For preparation of whole cell extracts, cells were incubated in RIPA buffer (50 mM Tris-HCl pH 7.5, 150 mM NaCl, 1% NP-40, 0,5% sodium deoxycholate, 0.1% SDS, 2 mM EDTA and 50mM NaF) supplemented with protease and phosphatase inhibitors, for 20 min on ice. Cells were scraped and centrifuged at 4 °C for 7 min at 17,000 x g to collect the supernatant containing the protein. The samples were applied to SDS-PAGE and analyzed by immunoblotting. Dilutions of primary antibodies used for immunoblotting were as follows: mouse monoclonal anti-FLAG M2 (Sigma-Aldrich), 1:2000; rabbit polyclonal anti-RtcB (Proteintech), 1:1000; for recognition of the human PTP1B, mouse monoclonal anti-PTP1B (3A7) (Santa Cruz Biotechnologies), 1:1000; for recognition of the mouse PTP1B, rabbit polyclonal anti-PTP1B (ab88481) (Abcam), 1:1000; rabbit polyclonal anti-IRE1α (B-12) (Santa Cruz Biotechnologies), 1:1000; mouse cocktail anti-pY-HRP (PY-7E1, PY20) (Invitrogen/Thermo-Fisher Scientific), 1:2000, goat polyclonal anti-Actin (I-19) (Santa Cruz Biotechnologies), 1:1000; mouse monoclonal anti-VCP (BD Transduction Laboratories), 1:1000; rabbit polyclonal anti-CNX/Calnexin (kindly provided by Dr. John Bergeron, McGill University, Montreal, QC, Canada), 1:1000. Polyclonal goat anti-mouse, goat anti-rabbit and rabbit anti-goat secondary antibodies conjugated to HRP (Dako-Agilent) were used at a dilution of 1:7000, 1:7000 and 1:3500, respectively. For the immunoprecipitation (IP), the cells were lysed in CHAPS buffer (30 mM Tris-HCl pH7.5, 150 mM NaCl and 1.5% CHAPS) supplemented with a cocktail of protease and phosphatase inhibitors. The cell lysis buffer was further supplemented with 1mM of the phosphatase inhibitor Na_3_VO_4_ when the immunoprecipitates were going to be tested for pTyr. The lysates were then incubated for 16 h at 4°C with the indicated IP antibody (1 μg Ab/1000 μg protein). After this, dynabeads protein G (Thermo-Fisher Scientific) were first washed with CHAPS lysis buffer and/or PBS, then mixed with the protein/Ab mixture, incubated at room temperature for 20 min or at 4°C for 40 min with gentle rotation and washed with CHAPS and/ or PBS. Finally, the beads were eluted with 1x Laemmli Sample Buffer (LSB), heated at 55-95°C for 5 min and loaded to SDS-PAGE. For the immunoblotting, the suitable primary and secondary antibodies were used.

### Generation of pY475-RtcB specific antibodies

These antibodies were generated by Biotem (France). Briefly, two rabbits were immunized with a KLH-conjugated MEEAPES(Phospho)YKNVTDVV peptide (4 immunizations). Sera were collected 42 days after immunization and specific antibodies subjected to affinity purification using MEEAPES(Phospho)YKNVTDVV-conjugated beads as affinity matrix. The bound material was eluted using acidic pH and then counter depleted for antibodies against the non-phosphorylated peptide using MEEAPESYKNVTDVV-conjugated beads. The flow-through was collected and further validated (**Fig. S3**).

### RT-PCR and RT-qPCR

Total RNA was extracted from cells using Trizol (Thermo-Fisher Scientific) according to the manufacturer’s instructions. cDNA was synthesized from the total RNA using the Maxima Reverse Transcriptase enzyme, random hexamer primers, dNTP mix and the Ribolock RNase inhibitor (Thermo-Fisher Scientific). PCR was performed on the template cDNA using Phusion High-Fidelity DNA Polymerase and dNTP mix (Thermo-Fisher Scientific). Quantitative PCR was alternatively performed for the cDNA using the SYBR® Premix Ex Taq™ (Tli RNase H Plus) (TAKARA-Clontech) using a QuantStudio5 system (Applied Biosystems). The primer sequences used for these experiments were synthesized by Eurogentec and are shown in **Table S4**.

### Cell death assay

HeLa cells stably expressing RtcB-Flag proteins were left untransfected or transfected with 10nM of a duplex siRNA sequence (**Table S4**; IDT, Integrated DNA Technologies) targeting the 3’-UTR of the RTCB mRNA. Forty-eight hours later the cells were treated with 0, 1, 2.5, 5, 7.5 and 10 mM DTT or 0, 0.5, 1 and 2 μM Doxorubicin or 0, 5, 10 and 20 μM Etoposide for 24 hours. The cells were then collected in tubes, along with the corresponding supernatants and PBS washes, and centrifuged at 1700 rpm or 750 xg for 5 min. The cell pellets were transferred in a round-bottom 96-well plate where 1x Annexin V buffer and Annexin V were added. After a 15 min-incubation at room temperature and a PBS wash, 2% FCS in PBS and 7-AAD were added to the samples. The latter were incubated at room temperature for 5 more min and then analysed by FACS (NovoCyte 3000).

### In vitro kinase assays

To perform an *in vitro* kinase assay, we used 0.36-0.5 μg of human recombinant His-tagged ABL (His-ABL1 or c-ABL: 126 kDa) from Carna Biosciences as the tyrosine kinase and 1 μg of human recombinant GST-tagged RtcB (GST-C22orf28: 81.6 kDa) (P01) from Abnova as the substrate. These were incubated in 1x kinase buffer (25 mM Tris-HCl pH 7.5, 10 mM MgCl_2_, 1 mM MnCl_2_, 0.5 mM DTT, 10 μM ATP, 0.1 mM Na_3_VO_4_ and 5mM β-glycerophosphate) in a total volume of 30 μl at 37°C for 30 min or at RT for 2 hours. Upon completion of the reaction, 5 x LSB was added to the reaction samples which were heated to 100°C and applied in a 6% SDS-PAGE.

### In vitro reconstitution of XBP1u splicing

To reconstitute XBP1u splicing *in vitro*, a vector encoding XBP1u downstream of T7 promoter was used as template for the splicing substrate (plasmids were a kind gift from Dr Fabio Martinon, Lausanne, Switzerland). The splicing substrate XBP1u mRNA was prepared using a RiboMAX™ Large Scale RNA Production Systems—T7 kit (Promega) at 37°C for 3 hours. *In vitro* transcripts were purified with RNeasy kit (Qiagen). Flag-RtcB WT and Flag-RtcB Y306F proteins were purified from HeLa cells stably expressing the two proteins, respectively. Cell pellets were lysed in a buffer containing 30 mM Tris-HCl, pH 7.5, 150 mM NaCl and 1.5% CHAPS supplemented with a cocktail of protease and phosphatase. Flag-tagged RtcB proteins were affinity-purified from the lysates with anti-Flag M2 affinity-gel (Sigma) followed by extensive washes. The human recombinant IRE1 cytoplasmic domain was purchased from Sino Biological. The reconstituted *in vitro* XBP1u splicing assay was carried out at 37°C for 2 hours in Kinase buffer (2 mM ATP, 2 mM GTP, 50 mMTris-HCl pH 7.4, 150 mM NaCl, 1 mM MgCl2, 1 mM MnCl2, 5 mM β-mercaptoethanol). One μg XBP1u RNA, 250 ng IRE1 protein and IP-purified Flag-RtcB on beads, were used for a 50-μl splicing reaction. The spliced products were column-purified with RNeasy kit (Qiagen) for RT-PCR analysis. The PCR products were resolved on a 3% agarose gel.

### Mass spectrometry analyses

Samples for the mass spectrometry analysis came from a scaled-up *in vitro* kinase reaction containing 0.9 μg of human recombinant His-c-ABL (His-ABL1 or c-ABL: 126 kDa) and 5 μg of human recombinant GST-RtcB in the same conditions as described in the above section. The difference is that the samples were ran on a pre-cast 8% polyacrylamide gel (Eurogentec), which was afterwards stained with 10% CBB and destained to be sent for mass spectrometry analysis. At the proteomics platform, the gel was re-stained with Page Blue Protein Staining Solution (Thermo-Fisher Scientific). In-gel digestion was then performed with trypsin (cleavage site = K/R) or lysC (cleavage site = K) and TiO_2_ purification (10% not purified as control) followed. For the MS analysis, the recovery volume was 15µl 0.1%TFA and the injection volume was 5µl. The instrument LTQ Velos and the PepMap 25cm or Orbitrap ELITE / C18 Accucore 50cm columns were used. One hour run MSA and HCD method was used. For the data processing, the software Proteome Discoverer 2.1 (Sequest HT / Percolator) was used with the thresholds of 1% FDR, ptmRS for site assignation. Data are available via ProteomeXchange with identifier PXD023433.

### Modeling of WT and phosphorylated RtcB

The starting structure for the molecular dynamics (MD) simulations was a homology model of the human RtcB protein (Nandy *et al*, 2017). For the modified versions of the protein regarding its tyrosine phosphorylation states, we edited the initial structure through YASARA (Yet Another Scientific Artificial Reality Application, v15.3.8) (Krieger & Vriend, 2014). For the MD simulations using Gromacs 5.1 (Abraham *et al*, 2015), preparation and generation of additional missing parameters to the AMBER ff14SB force field (Maier *et al*, 2015) was performed for each structure using the AmberTools17 package (Case *et al*, 2017). Since RtcB contains a Mn^2+^ ion in its active site, we employed the python-based metal center parameter builder MCPB.py (version 3.0) (Li & Merz, 2016) included in AmberTools17 for the parameterization of the metal site. In addition, separate pdb files were generated for the non-standard amino acid of phosphorylated tyrosine (named “TYP”) and GMP bound to H428 (named “HIG”), using standard protocols. The resulting topology and coordinate files from Amber were transformed to the respective Gromacs files in order to perform the necessary system preparations for the MD simulations. Each system was solvated with TIP3P water (Jorgensen *et al*, 1998) under periodic boundary conditions with the minimum distance between any atom in the solute and the edge of the periodic box being 10.0 Å. Na^+^/Cl^-^ counterions were added as appropriate for the neutralization of the system. Energy minimization was conducted until the force was less than 1000 kJ mol^-1^ nm^-1^, followed by a 100ps NVT and a 100ps NPT equilibration, and a final 200ns classical MD simulation for each system, using the Gromacs 5.1 package (Abraham *et al*, 2015). The temperature was kept at 300 K by the velocity rescaling thermostat (Bussi *et al*, 2009) with a coupling constant of 0.1 ps. In the NPT equilibration and MD simulation the pressure was kept at 1.0 bar using the Parrinello-Rahman barostat (Parrinello & Rahman, 1981) with a coupling time of 2.0 ps. The leap-frog algorithm (Van Gunsteren & Berendsen, 1988) was used with an integration time-step of 2 fs and the LINCS algorithm (Hess *et al*, 1997) was used to apply constraints on all bonds. UCSF Chimera (Pettersen *et al*, 2004) was used to visualize the processed systems. For quality control of our simulations we calculated the RMSF and RMSD of our systems. Post-MD analyses also included calculation of the surface solvent accessible area via Gromacs, pKa values and percentage of surface/buried area of the key tyrosine residues via the PROPKA 3.0 part of the PDB2PQR web server 2.0.0 (Dolinsky *et al*, 2004). To determine the relative positions of the tyrosines of interest and illustrate if and how much they are rotated, we set as a fixed point the Mn^2+^ ion in the active site of RtcB and calculated the dihedral angles consisting of this metal ion and the CA, CB and O atoms of the tyrosine residue using UCSF Chimera. For the representation of the dihedral angles, polar charts were created using the matlablib 2.1.2 in python (Droettboom *et al*, 2018). Furthermore, we calculated the docking score of the ligand GTP in the active site of the different RtcB systems (without GMP in the active site). For this purpose, among the programs of the Schrödinger Suites 2018-1 package, we used Protein Preparation Wizard (Madhavi Sastry *et al*, 2013) to prepare the protein structure, LigPrep to prepare the GTP ligand and Glide (Friesner *et al*, 2006) for receptor grid generation and XP ligand docking. The grid in which the ligand was allowed to dock was determined by analyzing the interactions between GMP and its surrounding amino acids in MOE (Molecular Operating Environment, 2016.01; Chemical Computing Group) (Molecular Operating Environment (MOE), 2016).

### Modeling of protein complexes

To explore the interaction of RtcB with the IRE1-XBP1 complex, we used a recent model of the human IRE1 tetramer (dimer of dimers) (Carlesso *et al*, 2020), and the homology model of human RtcB (Nandy *et al*, 2017). The RNAComposer modeling webserver (http://rnacomposer.ibch.poznan.pl) (Popenda *et al*, 2012) was used to predict the 3D structure of XBP1 mRNA based on the nucleotide sequence and its secondary structure (Peschek *et al*, 2015). The predicted model was refined and optimized using 3dRNA v.2.0 (http://biophy.hust.edu.cn/3dRNA) (Wang *et al*, 2019). The HADDOCK (Van Zundert *et al*, 2016) webserver was employed to predict the IRE1-tetramer/XBP1 complex. The cleavage site in both stem loops of XBP1 mRNA (G and C nucleotides) (Popenda *et al*, 2012) and active residues in the RNase domain of the IRE1 tetramer (*i.e.* His910 and Tyr892) (Sanches *et al*, 2014) were defined as interacting regions in HADDOCK. For the docking of RtcB and its pTyr306 phosphorylated variant to either the IRE1 tetramer or the IRE1 tetramer/XBP1 complex, the webserver HDOCK (Yan *et al*, 2017) was employed, using a blind-docking strategy. The Schrödinger package (Schrödinger Inc, release 2020-4) was employed to calculate the free energy of binding in IRE1-tetramer/XBP1 and IRE1-XBP1/RtcB complexes using the molecular mechanics generalized Born surface area (MM-GBSA) technique (Rastelli *et al*, 2010). For the c-Abl – RtcB interaction, we used the active form of Abl (PDB-ID: 2GQG). The ATP molecule was aligned inside protein using Chimera, based on the crystal structure of ATP bound to the structurally similar kinase Src (PDB-ID: 5XP7). The proteins were docked using PatchDock (Schneidman-Duhovny *et al*, 2005), with Y306 (RtcB) and ATP (c-Abl) chosen as interacting units. Following visual inspection of obtained complexes, the best aligned complex was prepared for MD simulations as outlined above. In order for the complex to not immediately dissociate, a set of seven position restrained equilibrations, with gradually reduced restraints in each, were applied. Finally, the system was subjected to a 200 ns MD simulation without position restraints.

### Bioinformatics tools

For the creation of Venn diagrams, we used the online tool http://bioinformatics.psb.ugent.be/webtools/Venn/. For the creation of gene networks and functional annotations, we used the online String database 11.0 (https://string-db.org/) (Jensen *et al*, 2009). Multiple sequence alignments were done using Clustal Omega 1.2.4 (Sievers *et al*, 2011). The PhosphoSitePlus database was used for the search of reported tyrosine phosphorylation sites on the proteins of interest (Hornbeck *et al*, 2015).

### Statistical analyses

Student’s t-test, one-way ANOVA and two-way ANOVA were applied for the statistical analyses depending on the experimental setting through the GraphPad Prism 7.0a software.

## Acknowledgements

We thank Dr Anna Reymer, Dr Antonio Carlesso, Dr Samuel Genheden and Johanna Hörberg for training on computational chemistry/biology (UGOT, Göteborg, Sweden), Dr Aeid Igbaria for critical reading of the manuscript. This work was funded by grants from Institut National du Cancer (INCa PLBIO), SIRIC-ILIAD/Cancéropôle Grand Ouest to EC, Fondation pour la Recherche Médicale (FRM, équipe labellisée 2018) to EC and RP; by EU H2020 MSCA ITN-675448 (TRAINERS) and MSCA RISE-734749 (INSPIRED) grants to EC and LAE. The Swedish Research Council (VR) and the Swedish National Infrastructure for Computing (SNIC) are gratefully acknowledged for funding and allocations of computing time at the C3SE and PDC supercomputing centers, respectively (LAE). AP is a Marie Curie early stage researcher funded by EU H2020 MSCA ITN- 675448 (TRAINERS). AM was funded by the Fondation ARC pour la recherche contre le cancer. SJM was funded by the Vinnova Seal-of-Excellence programme 2019-02205 (CaTheDRA).

## References

1. Abraham MJ, Murtola T, Schulz R, Páll S, Smith JC, Hess B & Lindah E (2015) GROMACS: High performance molecular simulations through multi-level parallelism from laptops to supercomputers. SoftwareX 1–2: 19–25

2. Almanza A, Carlesso A, Chintha C, Creedican S, Doultsinos D, Leuzzi B, Luís A, McCarthy N, Montibeller L, More S, et al (2018) Endoplasmic reticulum stress signalling - from basic mechanisms to clinical applications. FEBS J 286: 241–278

3. Belyy V, Zuazo-Gaztelu I, Alamban A, Ashkenazi A & Walter P (2021) Endoplasmic reticulum stress activates human IRE1α through reversible assembly of inactive dimers into small oligomers. bioRxiv: 2021.09.29.462487

4. Bian Y, Li L, Dong M, Liu X, Kaneko T, Cheng K, Liu H, Voss C, Cao X, Wang Y, et al (2016) Ultra-deep tyrosine phosphoproteomics enabled by a phosphotyrosine superbinder. Nat Chem Biol 12: 959–966

5. Blanchetot C, Chagnon M, Dube N, Halle M & Tremblay M (2005) Substrate-trapping techniques in the identification of cellular PTP targets. Methods 35: 44–53

6. Bussi G, Zykova-Timan T & Parrinello M (2009) Isothermal-isobaric molecular dynamics using stochastic velocity rescaling. J Chem Phys 130: 074101

7. Calfon M, Zeng H, Urano F, Till JH, Hubbard SR, Harding HP, Clark SG & Ron D (2002) IRE1 couples endoplasmic reticulum load to secretory capacity by processing the XBP-1 mRNA. Nature 415: 92–96

8. Carlesso A, Hörberg J, Reymer A & Eriksson LA (2020) New insights on human IRE1 tetramer structures based on molecular modeling. Sci Rep 10: 17490

9. Case DA, Cerutti DS, Cheatham TE, III, Darden TA, Duke RE, Giese TJ, Gohlke H, Goetz AW, Greene D, et al (2017) AMBER 2017 San Francisco

10. Dolinsky TJ, Nielsen JE, McCammon JA & Baker NA (2004) PDB2PQR: an automated pipeline for the setup of Poisson-Boltzmann electrostatics calculations. Nucleic Acids Res 32: W665–7

11. Droettboom M, Caswell TA, Hunter J, Firing E, Nielsen JH, Varoquaux N, Lee A, Andrade ES de, Root B, Stansby D, et al (2018) matplotlib/matplotlib v2.1.2.

12. Drogat B, Auguste P, Nguyen DT, Bouchecareilh M, Pineau R, Nalbantoglu J, Kaufman RJ, Chevet E, Bikfalvi A & Moenner M (2007) IRE1 Signaling Is Essential for Ischemia-Induced Vascular Endothelial Growth Factor-A Expression and Contributes to Angiogenesis and Tumor Growth In vivo. Cancer Res 67: 6700–6707

13. Dufey E, Bravo-San Pedro JM, Eggers C, González-Quiroz M, Urra H, Sagredo AI, Sepulveda D, Pihán P, Carreras-Sureda A, Hazari Y, et al (2020) Genotoxic stress triggers the activation of IRE1α-dependent RNA decay to modulate the DNA damage response. Nat Commun 11: 2401

14. Friesner RA, Murphy RB, Repasky MP, Frye LL, Greenwood JR, Halgren TA, Paul C. Sanschagrin A & Mainz DT (2006) Extra Precision Glide: Docking and Scoring Incorporating a Model of Hydrophobic Enclosure for Protein−Ligand Complexes. J Med Chem 49: 6177–96

15. Gu F, Nguyên DT, Stuible M, Dubé N, Tremblay ML & Chevet E (2004) Protein-tyrosine phosphatase 1B potentiates IRE1 signaling during endoplasmic reticulum stress. J Biol Chem 279: 49689–93

16. Van Gunsteren WF & Berendsen HJC (1988) A Leap-Frog Algorithm for Stochastic Dynamics. Mol Simul 1: 173–185

17. Han D, Lerner AG, Walle L Vande, Upton J-P, Xu W, Hagen A, Backes BJ, Oakes SA & Papa FR (2009) IRE1α Kinase Activation Modes Control Alternate Endoribonuclease Outputs to Determine Divergent Cell Fates. Cell 138: 562–575

18. Hess B, Bekker H, Berendsen HJC & Fraaije JGEM (1997) LINCS: A Linear Constraint Solver for Molecular Simulations. 18: 1463–1472

19. Hetz C & Glimcher LH (2009) Fine-Tuning of the Unfolded Protein Response: Assembling the IRE1α Interactome. Mol Cell 35: 551–561

20. Hetz C & Papa FR (2018) The Unfolded Protein Response and Cell Fate Control. Mol Cell 69: 169–181

21. Higa A, Taouji S, Lhomond S, Jensen D, Fernandez-Zapico ME, Simpson JC, Pasquet JM, Schekman R & Chevet E (2014) Endoplasmic Reticulum Stress-Activated Transcription Factor ATF6α Requires the Disulfide Isomerase PDIA5 To Modulate Chemoresistance. Mol Cell Biol 34: 1839–1849

22. Hollien J, Lin JH, Li H, Stevens N, Walter P & Weissman JS (2009) Regulated Ire1- dependent decay of messenger RNAs in mammalian cells. J Cell Biol 186: 323–331

23. Hollien J & Weissman JS (2006) Decay of endoplasmic reticulum-localized mRNAs during the unfolded protein response. Science 313: 104–107

24. Hornbeck P V, Zhang B, Murray B, Kornhauser JM, Latham V & Skrzypek E (2015) PhosphoSitePlus, 2014: mutations, PTMs and recalibrations. Nucleic Acids Res 43: D512–20

25. Iwawaki T, Hosoda A, Okuda T, Kamigori Y, Nomura-Furuwatari C, Kimata Y, Tsuru A & Kohno K (2001) Translational control by the ER transmembrane kinase/ribonuclease IRE1 under ER stress. Nat Cell Biol 3: 158–165

26. Jensen LJ, Kuhn M, Stark M, Chaffron S, Creevey C, Muller J, Doerks T, Julien P, Roth A, Simonovic M, et al (2009) STRING 8--a global view on proteins and their functional interactions in 630 organisms. Nucleic Acids Res 37: D412–6

27. Jia Z, Barford D, Flint AJ & Tonks NK (1995) Structural Basis for Phosphotyrosine Peptide Recognition by Protein Tyrosine Phosphatase 1 B. Science 268: 1754–8

28. Jorgensen WL, Chandrasekhar J, Madura JD, Impey RW & Klein ML (1998) Comparison of simple potential functions for simulating liquid water. J Chem Phys 79: 926

29. Jurkin J, Henkel T, Nielsen AF, Minnich M, Popow J, Kaufmann T, Heindl K, Hoffmann T, Busslinger M & Martinez J (2014) The mammalian tRNA ligase complex mediates splicing of XBP1 mRNA and controls antibody secretion in plasma cells. EMBO J 33: 2922–36

30. Karagöz GE, Acosta-Alvear D, Nguyen HT, Lee CP, Chu F & Walter P (2017) An unfolded protein-induced conformational switch activates mammalian IRE1. Elife 6: e30700

31. Kosmaczewski SG, Edwards TJ, Han SM, Eckwahl MJ, Meyer BI, Peach S, Hesselberth JR, Wolin SL & Hammarlund M (2014) The RtcB RNA ligase is an essential component of the metazoan unfolded protein response. EMBO Rep 15: 1278–85

32. Krieger E & Vriend G (2014) YASARA View—molecular graphics for all devices—from smartphones to workstations. Bioinformatics 30: 2981

33. Krishnan N, Fu C, Pappin DJ & Tonks NK (2011) H2S-Induced Sulfhydration of the Phosphatase PTP1B and Its Role in the Endoplasmic Reticulum Stress Response. Sci Signal 4: ra86

34. Lee K, Tirasophon W, Shen X, Michalak M, Prywes R, Okada T, Yoshida H, Mori K & Kaufman RJ (2002) IRE1-mediated unconventional mRNA splicing and S2P-mediated ATF6 cleavage merge to regulate XBP1 in signaling the unfolded protein response. Genes Dev 16: 452–466

35. Lerner AG, Upton J-P, Praveen PVK, Ghosh R, Nakagawa Y, Igbaria A, Shen S, Nguyen V, Backes BJ, Heiman M, et al (2012) IRE1α Induces Thioredoxin-Interacting Protein to Activate the NLRP3 Inflammasome and Promote Programmed Cell Death under Irremediable ER Stress. Cell Metab 16: 250–264

36. Lhomond S, Avril T, Dejeans N, Voutetakis K, Doultsinos D, McMahon M, Pineau R, Obacz J, Papadodima O, Jouan F, et al (2018) Dual IRE1 RNase functions dictate glioblastoma development. EMBO Mol Med: e7929

37. Li P & Merz KM (2016) MCPB.py: A Python Based Metal Center Parameter Builder. J Chem Inf Model 56: 599–604

38. Li W, Crotty K, Garrido Ruiz D, Voorhies M, Rivera C, Sil A, Mullins RD, Jacobson MP, Peschek J & Walter P (2021) Protomer alignment modulates specificity of RNA substrate recognition by Ire1. Elife 10: e67425

39. Lu Y, Liang FX & Wang X (2014) A Synthetic Biology Approach Identifies the Mammalian UPR RNA Ligase RtcB. Mol Cell 55: 758–770

40. Madhavi Sastry G, Adzhigirey M, Day T, Annabhimoju R & Sherman W (2013) Protein and ligand preparation: parameters, protocols, and influence on virtual screening enrichments. J Comput Aided Mol Des 27: 221–234

41. Maier JA, Martinez C, Kasavajhala K, Wickstrom L, Hauser KE & Simmerling C (2015) ff14SB: Improving the Accuracy of Protein Side Chain and Backbone Parameters from ff99SB. J Chem Theory Comput 11: 3696–3713

42. Maurel M, Chevet E, Tavernier J & Gerlo S (2014) Getting RIDD of RNA: IRE1 in cell fate regulation. Trends Biochem Sci 39: 245–254

43. McGrath EP, Centonze FG, Chevet E, Avril T & Lafont E (2021) Death sentence: The tale of a fallen endoplasmic reticulum. Biochim Biophys acta Mol cell Res 1868: 119001

44. Molecular Operating Environment (MOE) 2013.08 (2016) Molecular Operating Environment (MOE), 2013.08; Chemical Computing Group Inc., 1010 Sherbooke St. West, Suite #910, Montreal, QC, Canada, H3A 2R7. Mol Oper Environ (MOE), 201308; Chem Comput Gr Inc, 1010 Sherbooke St West, Suite #910, Montr QC, Canada, H3A 2R7, 2013

45. Morita S, Villalta SA, Feldman HC, Register AC, Rosenthal W, Hoffmann-Petersen IT, Mehdizadeh M, Ghosh R, Wang L, Colon-Negron K, et al (2017) Targeting ABL- IRE1α Signaling Spares ER-Stressed Pancreatic β Cells to Reverse Autoimmune Diabetes. Cell Metab 25: 1207

46. Nandy A, Saenz-Méndez P, Gorman AM, Samali A & Eriksson LA (2017) Homology model of the human tRNA splicing ligase RtcB. Proteins Struct Funct Bioinforma 85: 1983–1993

47. Parrinello M & Rahman A (1981) Polymorphic transitions in single crystals: A new molecular dynamics method. J Appl Phys 52: 7182–7190

48. Peschek J, Acosta-Alvear D, Mendez AS & Walter P (2015) A conformational RNA zipper promotes intron ejection during non-conventional XBP1 mRNA splicing. EMBO Rep 16: 1688

49. Pettersen EF, Goddard TD, Huang CC, Couch GS, Greenblatt DM, Meng EC & Ferrin TE (2004) UCSF Chimera--A visualization system for exploratory research and analysis. J Comput Chem 25: 1605–1612

50. Popenda M, Szachniuk M, Antczak M, Purzycka KJ, Lukasiak P, Bartol N, Blazewicz J & Adamiak RW (2012) Automated 3D structure composition for large RNAs. Nucleic Acids Res 40: e112

51. Popow J, Jurkin J, Schleiffer A & Martinez J (2014) Analysis of orthologous groups reveals archease and DDX1 as tRNA splicing factors. Nature 511: 104–107

52. Rastelli G, Rio A Del, Degliesposti G & Sgobba M (2010) Fast and accurate predictions of binding free energies using MM-PBSA and MM-GBSA. J Comput Chem 31: 797–810

53. Ray A, Zhang S, Rentas C, Caldwell KA & Caldwell GA (2014) RTCB-1 mediates neuroprotection via XBP-1 mRNA splicing in the unfolded protein response pathway. J Neurosci 34: 16076–85

54. Sanches M, Duffy NM, Talukdar M, Thevakumaran N, Chiovitti D, Canny MD, Lee K, Kurinov I, Uehling D, Al-Awar R, et al (2014) Structure and mechanism of action of the hydroxy-aryl-aldehyde class of IRE1 endoribonuclease inhibitors. Nat Commun 5: 4202

55. Schneidman-Duhovny D, Inbar Y, Nussinov R & Wolfson HJ (2005) PatchDock and SymmDock: servers for rigid and symmetric docking. Nucleic Acids Res 33: W363– W367

56. Sepulveda D, Rojas-Rivera D, Rodríguez DA, Groenendyk J, Köhler A, Lebeaupin C, Ito S, Urra H, Carreras-Sureda A, Hazari Y, et al (2018) Interactome Screening Identifies the ER Luminal Chaperone Hsp47 as a Regulator of the Unfolded Protein Response Transducer IRE1α. Mol Cell 69: 238–252.e7

57. Sievers F, Wilm A, Dineen D, Gibson TJ, Karplus K, Li W, Lopez R, McWilliam H, Remmert M, Söding J, et al (2011) Fast, scalable generation of high-quality protein multiple sequence alignments using Clustal Omega. Mol Syst Biol 7: 539

58. Spiotto MT, Banh A, Papandreou I, Cao H, Galvez MG, Gurtner GC, Denko NC, Le QT & Koong AC (2010) Imaging the unfolded protein response in primary tumors reveals microenvironments with metabolic variations that predict tumor growth. Cancer Res 70: 78–88

59. Tam AB, Koong AC & Niwa M (2014) Ire1 has distinct catalytic mechanisms for XBP1/HAC1 splicing and RIDD. Cell Rep 9: 850–8

60. Le Thomas A, Ferri E, Marsters S, Harnoss JM, Modrusan Z, Li W, Rudolph J, Wang W, Wu TD, Walter P, et al (2021) Noncanonical mRNA decay by the endoplasmic-reticulum stress sensor IRE1α promotes cancer-cell survival. bioRxiv: 2021.03.16.435520

61. Tsai C-F, Wang Y-T, Yen H-Y, Tsou C-C, Ku W-C, Lin P-Y, Chen H-Y, Nesvizhskii AI, Ishihama Y & Chen Y-J (2015) Large-scale determination of absolute phosphorylation stoichiometries in human cells by motif-targeting quantitative proteomics. Nat Commun 6: 6622

62. Tsai Y-L, Ha DP, Zhao H, Carlos AJ, Wei S, Pun TK, Wu K, Zandi E, Kelly K & Lee AS (2018) Endoplasmic reticulum stress activates SRC, relocating chaperones to the cell surface where GRP78/CD109 blocks TGF-β signaling. Proc Natl Acad Sci 115: 201714866

63. Upton J-P, Wang L, Han D, Wang ES, Huskey NE, Lim L, Truitt M, McManus MT, Ruggero D, Goga A, et al (2012) IRE1α cleaves select microRNAs during ER stress to derepress translation of proapoptotic Caspase-2. Science 338: 818–22

64. Urano F, Wang X-Z, Bertolotti A, Zhang Y, Chung P, Harding HP & Ron D (2000) Coupling of Stress in the Endoplasmic Reticulum to Activation of JNK Protein Kinases by Transmembrane Protein Kinase IRE1. Science 287: 664–666

65. Wang J, Wang J, Huang Y & Xiao Y (2019) 3dRNA v2.0: An Updated Web Server for RNA 3D Structure Prediction. Int J Mol Sci 20

66. Yan Y, Zhang D, Zhou P, Li B & Huang S-Y (2017) HDOCK: a web server for protein– protein and protein–DNA/RNA docking based on a hybrid strategy. Nucleic Acids Res 45: W365–W373

67. Yang Z, Zhang J, Jiang D, Khatri P, Solow-Cordero DE, Toesca DAS, Koumenis C, Denko NC, Giaccia AJ, Le Q-T, et al (2018) A Human Genome-Wide RNAi Screen Reveals Diverse Modulators that Mediate IRE1α–XBP1 Activation. Mol Cancer Res 16: 745–753

68. Zhang Y-L, Yao Z-J, Sarmiento M, Wu L, Burke TR & Zhang Z-Y (2000) Thermodynamic Study of Ligand Binding to Protein-tyrosine Phosphatase 1B and Its Substrate-trapping Mutants. 275: 34205–12

69. Van Zundert GCP, Rodrigues JPGLM, Trellet M, Schmitz C, Kastritis PL, Karaca E, Melquiond ASJ, Van Dijk M, De Vries SJ & Bonvin AMJJ (2016) The HADDOCK2.2 Web Server: User-Friendly Integrative Modeling of Biomolecular Complexes. J Mol Biol 428: 720–725

