## Supplementary material for "Stress-induced tyrosine phosphorylation of RtcB modulates IRE1 activity and signaling outputs": Suppl material

**Papaioannou et al.**

**Supplemental material**

**TABLES S1-S4**

**FIGURES S1-S10**

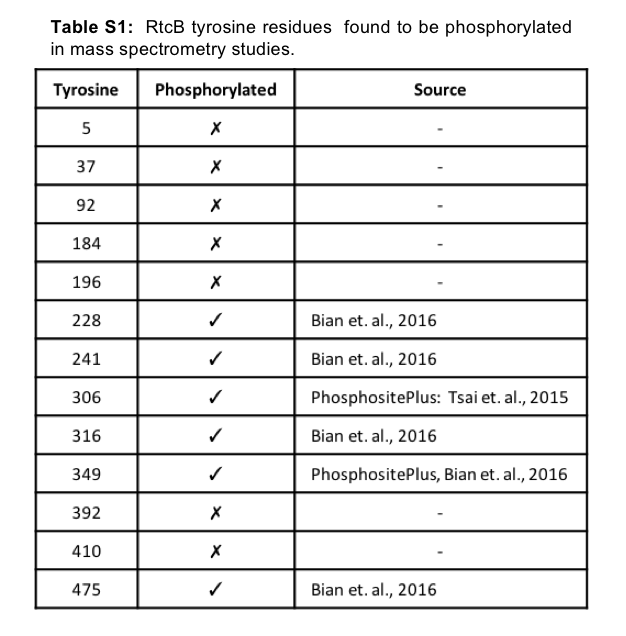

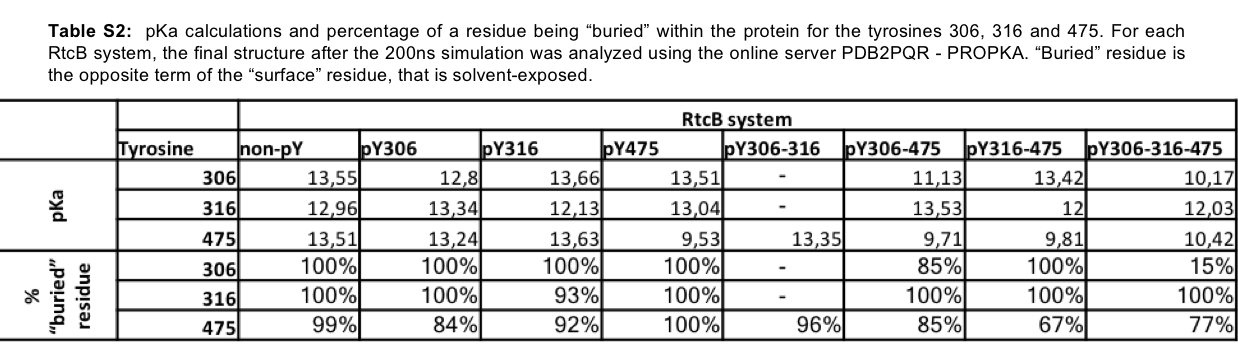

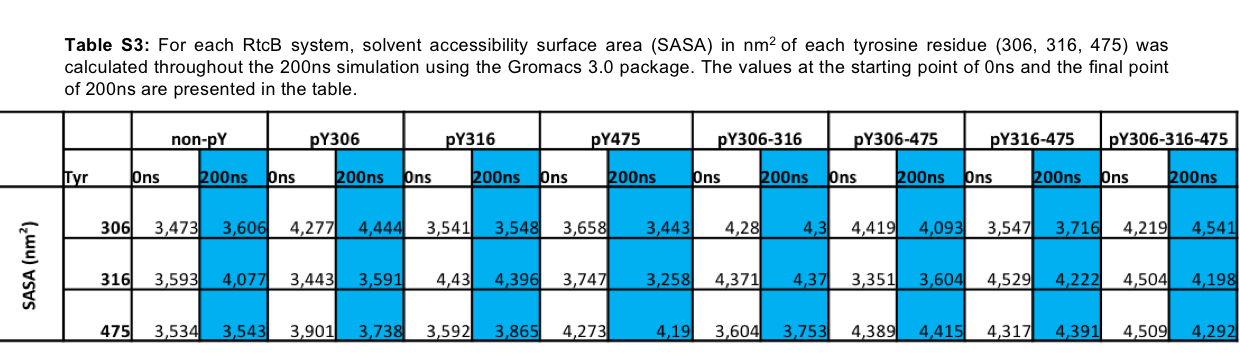

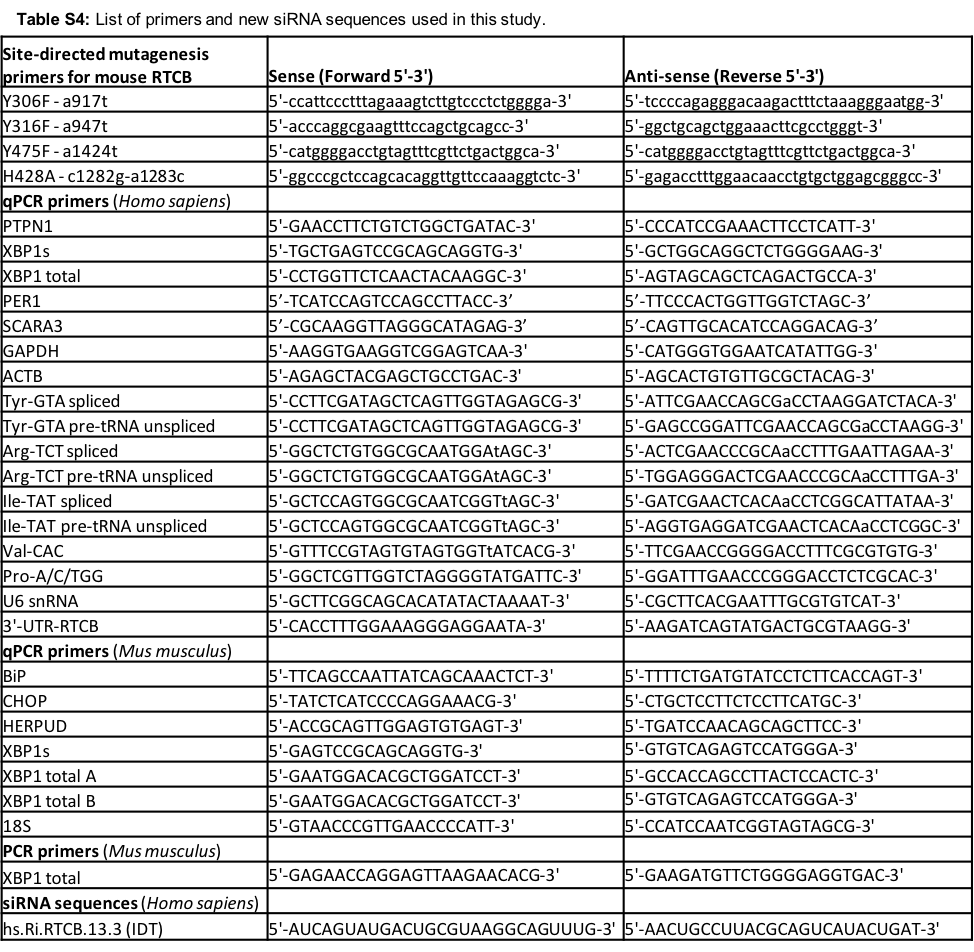

**Figure S1.** (**A**) The XBP1s-luciferase reporter described in detail in “Materials and Methods”. (**B**) An in vitro IRE1α-mediated cleavage assay (described in “Materials and Methods” and in [23]). (**C**) Gene ontology (GO) annotation of the gene network shown in **Fig 1D**. (**D**) Scheme of the regions that XBP1s and XBP1 total primers amplify on the XBP1 unspliced or spliced mRNA (used in Fig. 1E). (**E-G**) mRNA expression levels of BiP (E), CHOP (F) and HERPUD (G) in PTP1B^+/+^ or PTP1B^-/-^ MEFs untreated or treated with 10μg/ml Tun for 3 and 6 hours. (**H**) mRNA expression levels of XBP1s/total, BiP, CHOP and HERPUD in PTP1B^+/+^ or PTP1B^-/-^ MEFs under basal conditions. (**I**) PTP1B mRNA expression levels in U87 cells expressing or not a dominant negative form of IRE1α during Tun treatment (5μg/ml) with a 2-hour Actinomycin D (5μg/ml) pre-treatment. EV: empty vector, IRE1 DN: dominant negative form (cytosolic-deficient) IRE1α. (**J**) PTP1B protein level in U87 cells during Tun treatment (5μg/ml) in different time points. Data values of (E-H) come from one experiment, while data values of (I-J) are the mean ± SEM of n≥3 independent experiments (***p < 0.001, ****<0.0001). Error bars for (I) are not depicted.

**Figure S2.** (**A**) Multiple sequence alignment for the tRNA-splicing ligase RtcB homolog of *Homo sapiens, Mus musculus, Drosophila melanogaster, Ceanorhabditis elegans* and *Danio rerio*, representative organisms of the animal kingdom. Only the parts of the sequences including the tyrosines that can be phosphorylated in the human RtcB protein and are known from MS studies are shown. (**B**) Samples as represented in Fig. 2A were treated or not with 15μM bpV(phen) for 2 hours and their protein lysates were analyzed for Flag, RtcB, pY-HRP and VCP. (**C**) HEK293T cells were treated with 15μM bpV(phen) for 2 hours and their protein lysates were immunoprecipitated for the endogenous RtcB protein using an anti-RtcB antibody. The resulting sample was probed for pY and the membrane was re-probed for RtcB. (**D**) In vitro kinase assay containing recombinant human proteins of c-Abl (His-c-Abl) and RtcB (GST-RtcB) in the combinations seen in the three different lanes of the blots. The completed reactions were ran on a polyacrylamide gel and the membrane was immunoblotted for phosphotyrosine and then re-probed for RtcB. The arrowhead points to the GST-RtcB protein and the asterisk denotes the auto-phosphorylated His-c-Abl. The depicted western blots are representative of independent experiments.

**Figure S3.** (**A**) Schematic representation of the experimental pipeline for generating anti pY475-RtcB antibodies. (**B**) Western blot analysis of lysates from HEK293T cells transfected with an empty vector (CTL) or an expression plasmid containing Flag-RtcB-WT or Flag-RtcB-Y475F using 2 affinity purified antibodies against pY475 (#207, #234). (**C**) Western blot analysis of anti-Flag immunoprecipitates from HEK293T cells transfected with an empty vector (CTL) or an expression plasmid containing Flag-RtcB-WT or Flag-RtcB-Y475F using the #207 affinity purified antibodies against pY475.

**Figure S4**. (**A**) Zoom-in into the active site of the unphosphorylated RtcB system after 200 ns MD simulation. (**B**) The systems with RtcB phosphorylated on Tyr 306, 316, 475, 306 and 316, 306 and 475, 316 and 475 are shown after the 200 ns MD simulations.

**Figure S5.** (**A**) Zoom-in into the active site containing GMP bound to His 428 of the unphosphorylated RtcB system after 200 ns MD simulation. (**B**) All RtcB systems containing GMP bound to the nitrogen of the histidine 428 in their active site, after 200 ns MD simulations. The systems represent the state after the attack of GTP, whereby RtcB is ready to proceed with the catalytic ligation of the spliced ends of XBP1 mRNA.

**Figure S6.** (**A**) HEK293T cells transfected with the FLAG-tagged phosphoablating mutants of RtcB (Y306F, Y316F and Y475F) were treated 24 hours post-transfection with 1ug/ml TM for 6 hours or transfected with c-ABL siRNA for 2 days. Immunoprecipitation was done in the cell lysates using Flag antibody and the immunoprecipitates were first immunoblotted for phosphotyrosine and then re-probed with Flag Ab. The levels of phosphorylated RtcB was then normalized to RtcB protein levels. Data shown in the graph correspond to the mean ± SEM of n=3 independent experiments. (**B**) HEK cells were left untransfected (CTL) or transfected with 1μg of each one of the Y-to-F RtcB-Flag mutant plasmids and with 1μg of the PTP1B C215S mutant plasmid. Immunoprecipitation was done in the cell lysates with the PTP1B antibody and the immunoprecipitates were immunoblotted with anti-RtcB antibodies and anti-PTP1B antibodies. Input lysates were immunoblotted with anti-RtcB, anti-Flag, anti-PTP1B and anti-Calnexin (CNX, used as loading control). (**C**) RtcB protein levels in the immunoprecipitated fractions of (A) after normalization to the immunoprecipitated PTP1B. Data values are the mean ± SEM of n=3 independent experiments. Unpaired t test was applied for the statistical analyses (*p<0.05, **p<0.01). Double: Y306F/Y316F; Triple: Y306F/Y316F/Y475F.

**Figure S7. (A)** HeLa cells expressing or not various Flag tagged versions of RtcB (WT, single mutants, double or triple mutants) were treated for 0, 1, 2, 4, 8h with 10μg/ml cycloheximide. Cell lysates were harvested and the expression of Flag-RtcB, endogenous RtcB or VCP evaluated using western blot with the corresponding antibodies. (**B-D**) Quantification of Fig. 4D for the different HeLa cell lines. Data values are the mean ± SEM of n=3 independent experiments. Error bars for (A-C) are not depicted. (**E**) HeLa lines were treated with 5μg/ml Tun at different time points. The graph shows the mRNA levels of PER1 and SCARA3 after normalization to the untreated samples (0h time-point). Data values come from one experiment. (**F**) *In vitro* mediated cleavage of XBP1 mRNA. Uncleavable XBP1 mRNA or cleavable XBP1 mRNA were incubated with increasing amounts of IRE1 (100-1000ng) in the conditions defined in Materials and Methods. The reaction products were then retrotranscribed, analyzed by PCR using primers recognizing the XBP1 mRNA and resolved on agarose gels by electrophoresis before being visualized. These experiments allowed us to determine the minimal concentration of IRE1 needed to specifically cleave XBP1 mRNA at splicing sites.

**Figure S8.** (**A**) PTP1B^+/+^ (wt) and PTP1B^-/-^ (KO) MEFs were treated with 10, 50 or 100ng/ml Tun for 0, 2, 4, 8, 16 and 24 hours. Their cDNA was analyzed by PCR using primers recognizing the XBP1 mRNA. (**B**) Quantification of the gels in (A). (**C**) PTP1B^+/+^ (wt) and PTP1B^-/-^ (KO) MEFs untransfected (CTL) or transfected with 2μg of wt or Y306F RtcB-Flag plasmids were treated 24 hours post-transfection with 10μg/ml Tun for 6 hours. The resulting RNA was analyzed using RT-qPCR for XBP1s and XBP1 total, thereby allowing XBP1 mRNA splicing using the ratio of both values. XBP1 mRNA splicing is represented as the ratio of the XBP1 splicing in the WT towards WT MEFs that equals 1 for every condition, and this ratio in the KO towards the WT cells for each condition. (**D**) Protein samples of the untreated cells in Fig. 6F were analyzed for the expression of the RtcB-Flag protein, PTP1B and VCP (loading control) using immunoblotting. (**E**) Protein complex after docking of c-Abl to the Y306 site of RtcB and 200ns MD simulation. Ribbon-like structures are shown. (**F**) Zoom-in on the interacting residues D325 and D444 of c-Abl (in green) and K279, R283 and K357 of RtcB (in red). (**G**) cDNA samples from Fig. 7A were analyzed for endogenous RTCB mRNA levels using suitable primers. U6 snRNA was used as the loading control. (**H**) Protein samples from Fig. 7A were analyzed for expression of the RtcB-Flag protein and CNX which served as a loading control. The blots shown here are representative of three or more independent experiments. Data represented are the mean ± SEM of n≥3 independent experiments. Unpaired t test was applied for the statistical analyses (ns: non-significant, *p < 0.05, ****p < 0.0001). Error bars for (B) are not depicted.

**Figure S9**. HeLa lines stably expressing wt or Y306F RtcB-Flag were left untransfected or transfected with siRNA sequences against the endogenous RTCB mRNA. 48 hours post-transfection they were treated with 0, 0.5, 1 and 2 uM Doxorubicin for 24 hours. The resulting samples were then analyzed by FACS for cell necrosis (Annexin-v) through 7-AAD staining. Data values are the mean ± SEM of n = 4 independent experiments in two-way ANOVA and post-Tukey multiple comparisons test (ns: non-significant, *p < 0.05, ***p < 0.001, ****p < 0.0001). Experiments presented in Fig. S9 and S10 were carried out simultaneously and therefore controls are identical in both figures.

**Figure S10**. HeLa lines stably expressing wt or Y306F RtcB-Flag were left untransfected or transfected with siRNA sequences against the endogenous RTCB mRNA. 48 hours post-transfection they were treated with 0, 5, 10 and 20 uM Etoposide for 24 hours. The resulting samples were then analyzed by FACS for cell necrosis (Annexin-V) through 7-AAD staining. Data values are the mean ± SEM of n = 4 independent experiments in two-way ANOVA and post-Tukey multiple comparisons test (ns: non-significant, *p < 0.05, ***p < 0.001, ****p < 0.0001). Experiments presented in Fig. S9 and S10 were carried out simultaneously and therefore controls are identical in both figures.

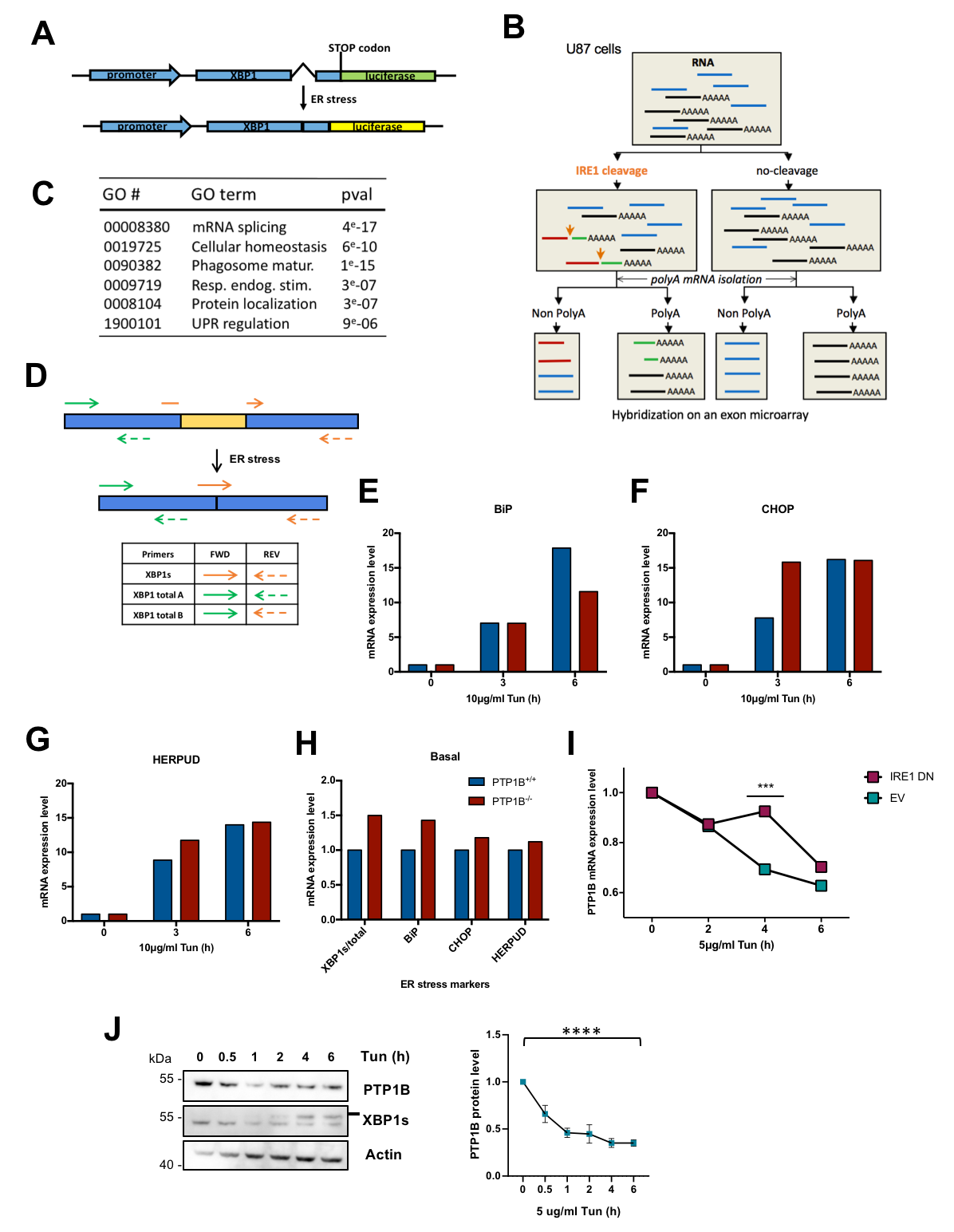

**FIGURE S1**

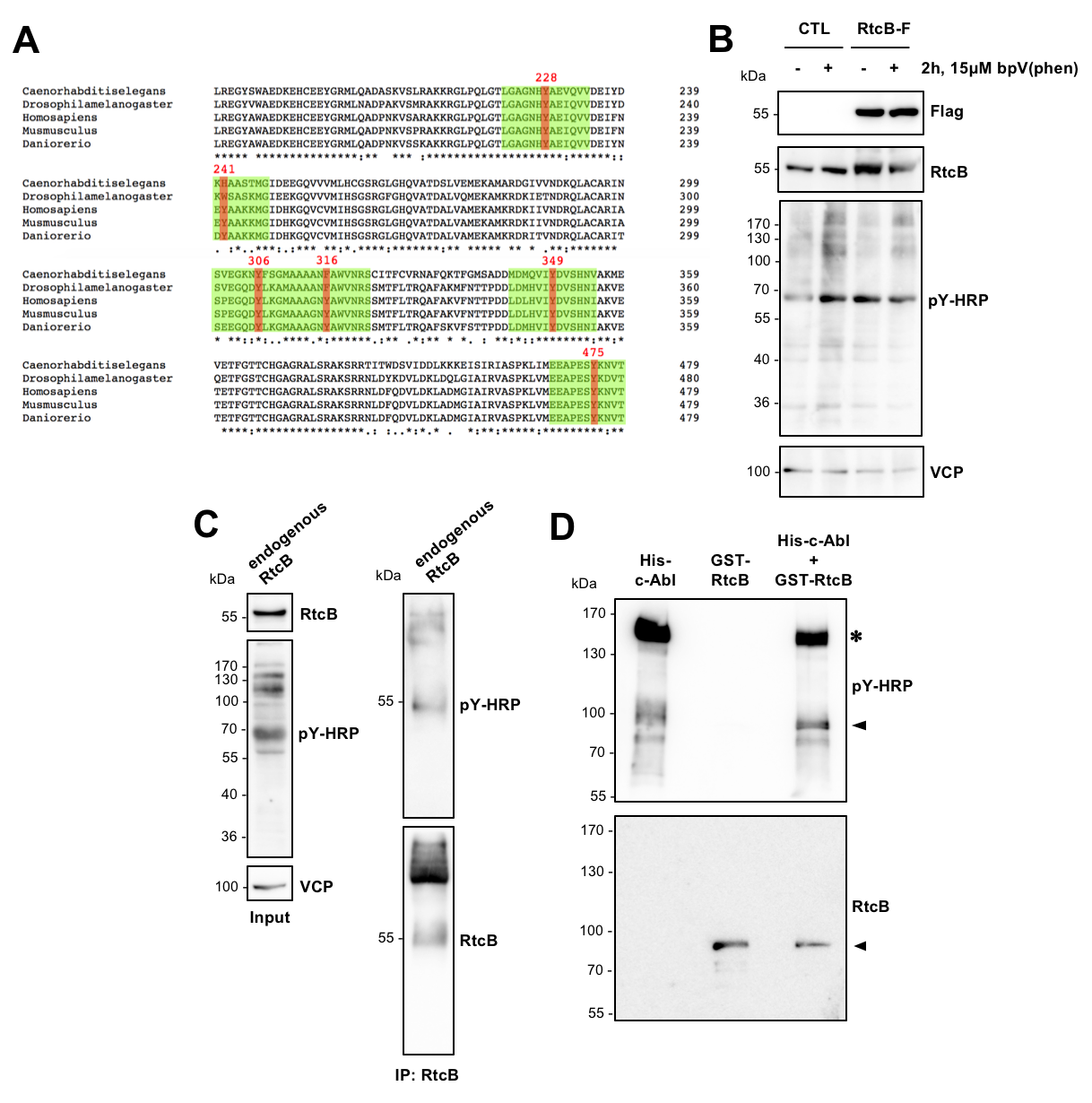

**FIGURE S2**

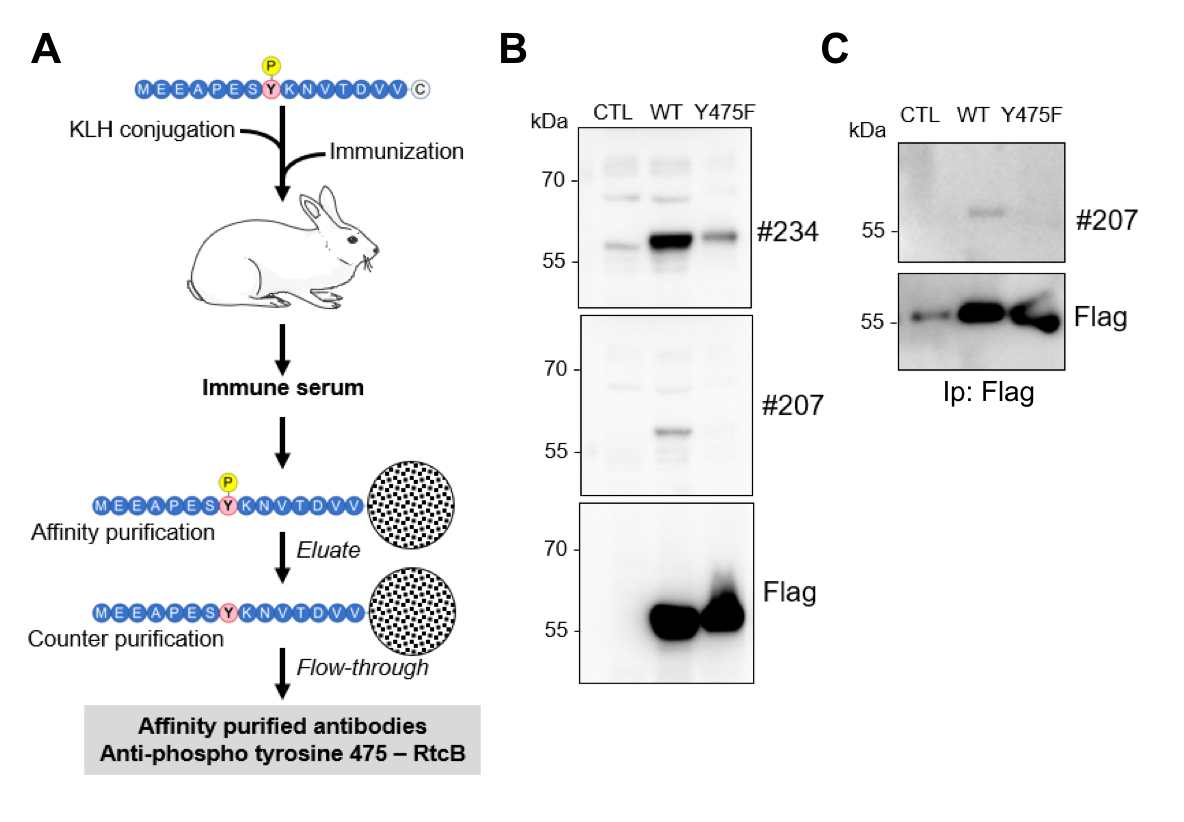

**FIGURE S3**

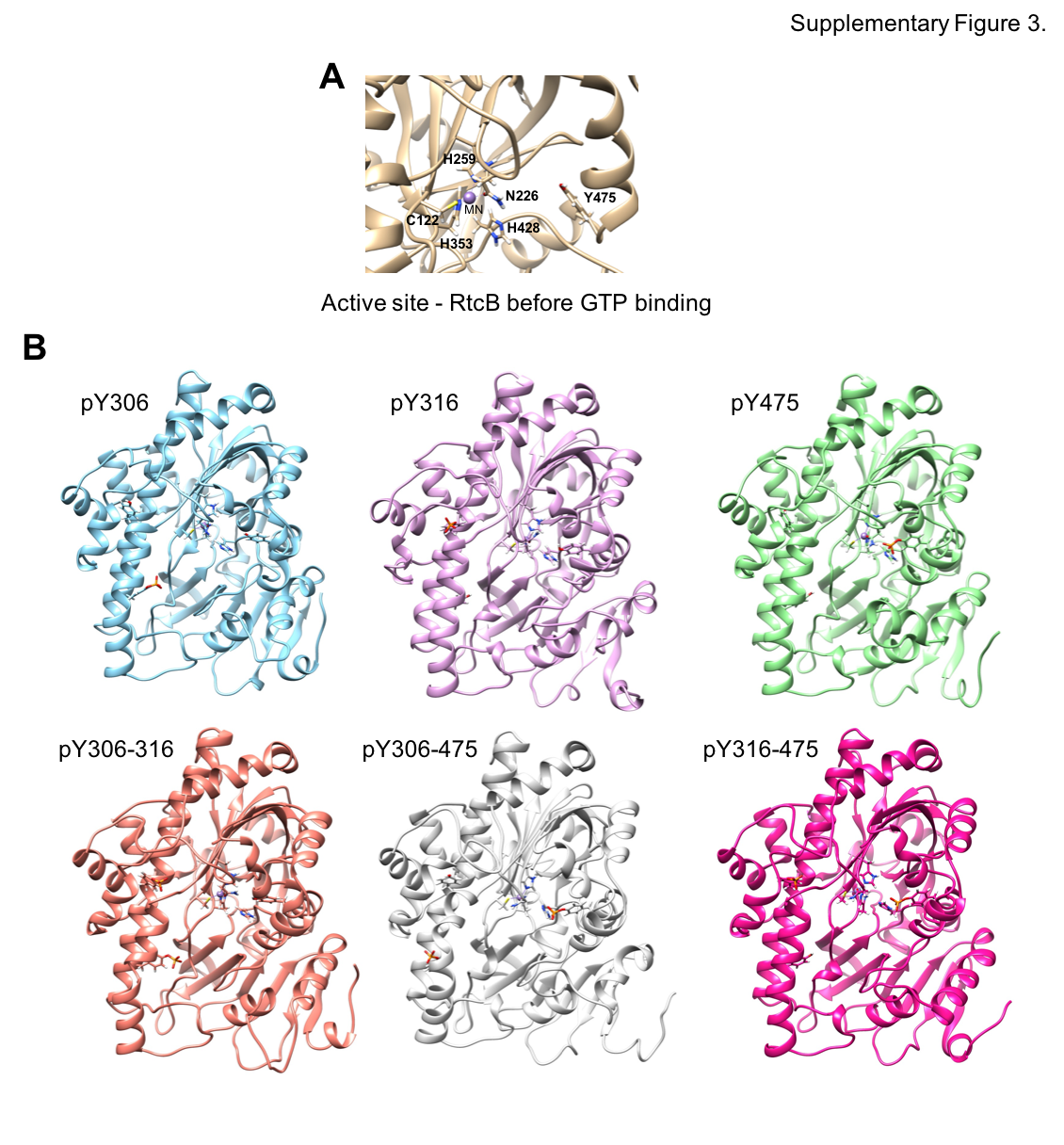

**FIGURE S4**

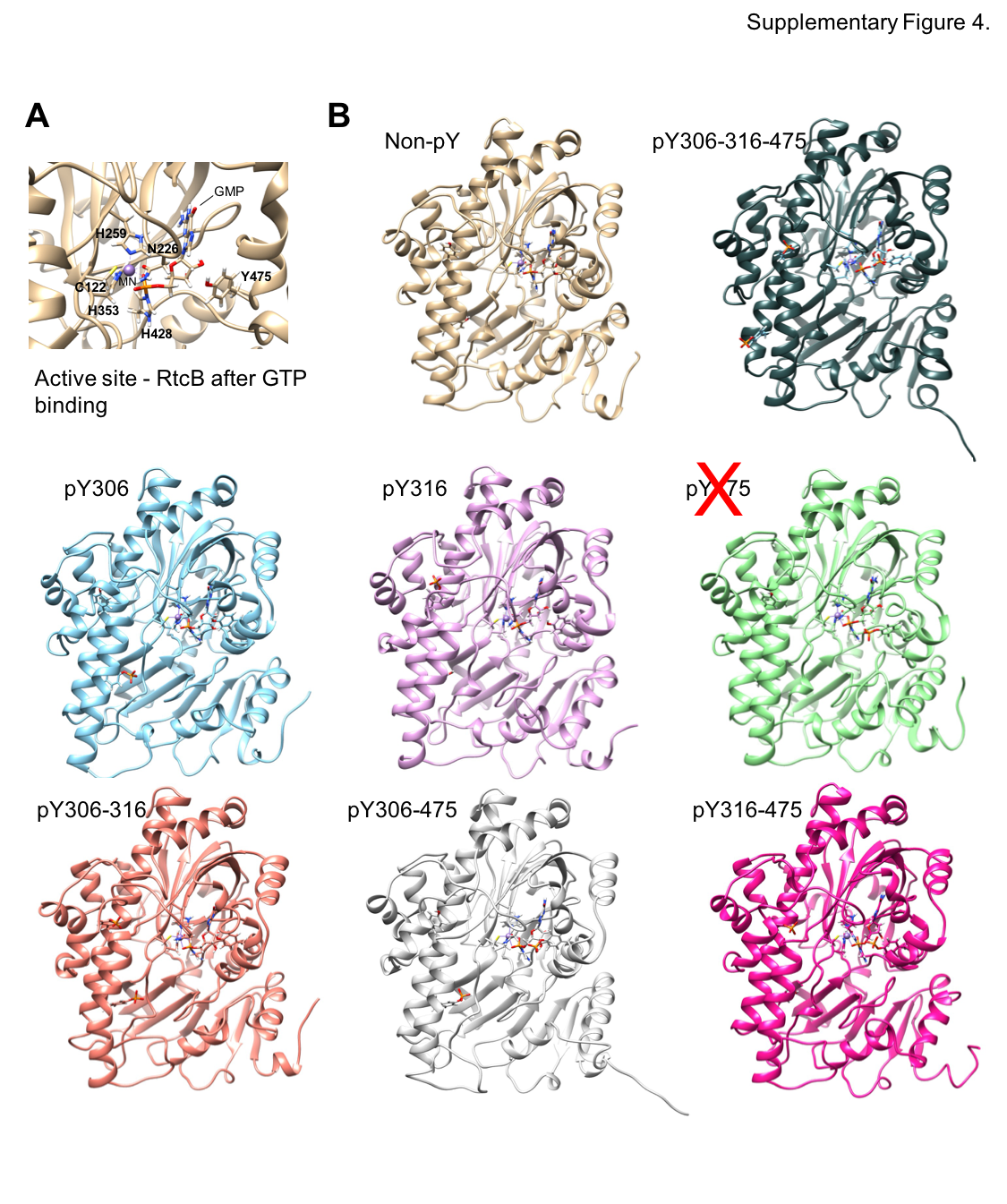

**FIGURE S5**

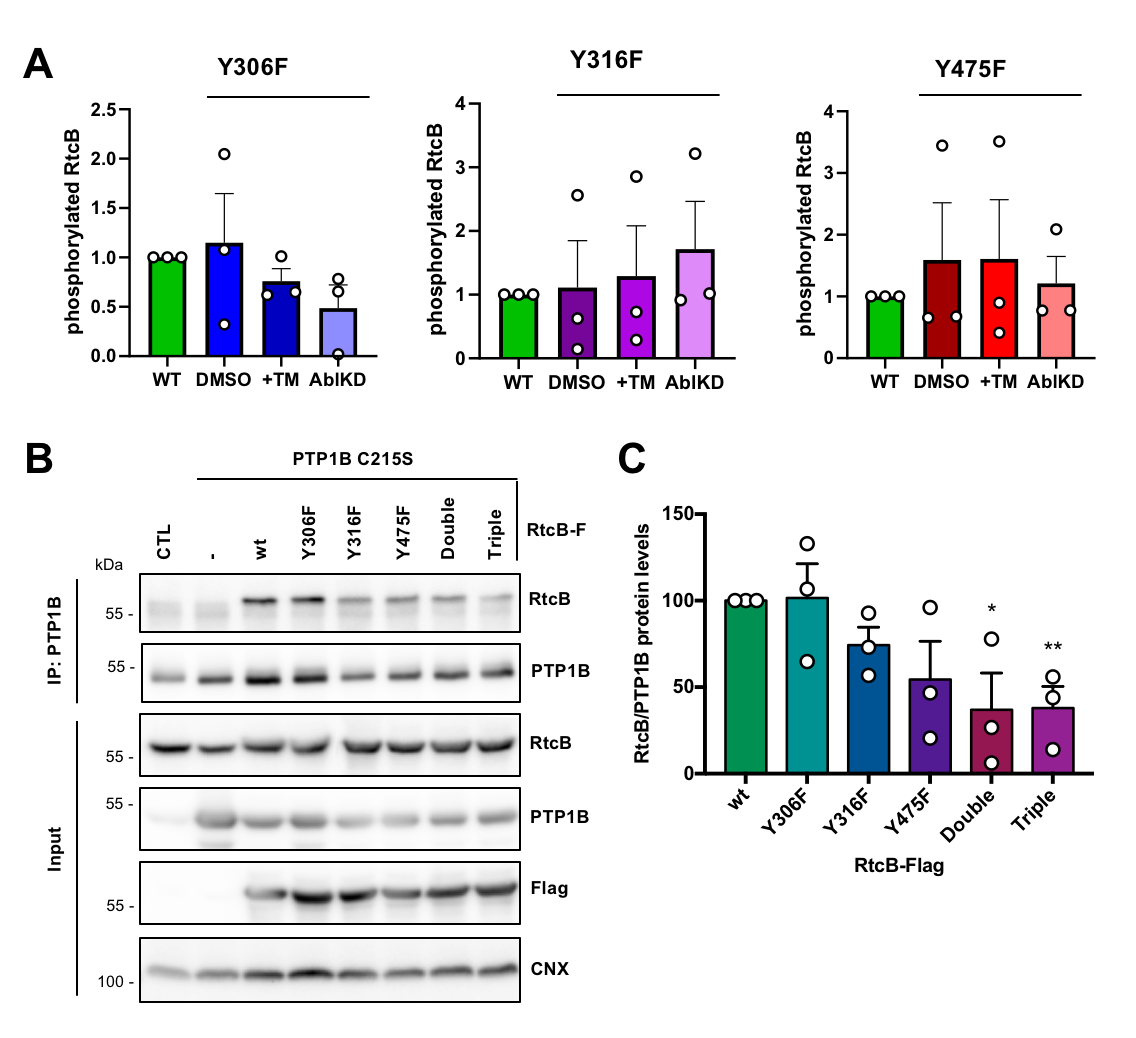

**FIGURE S6**

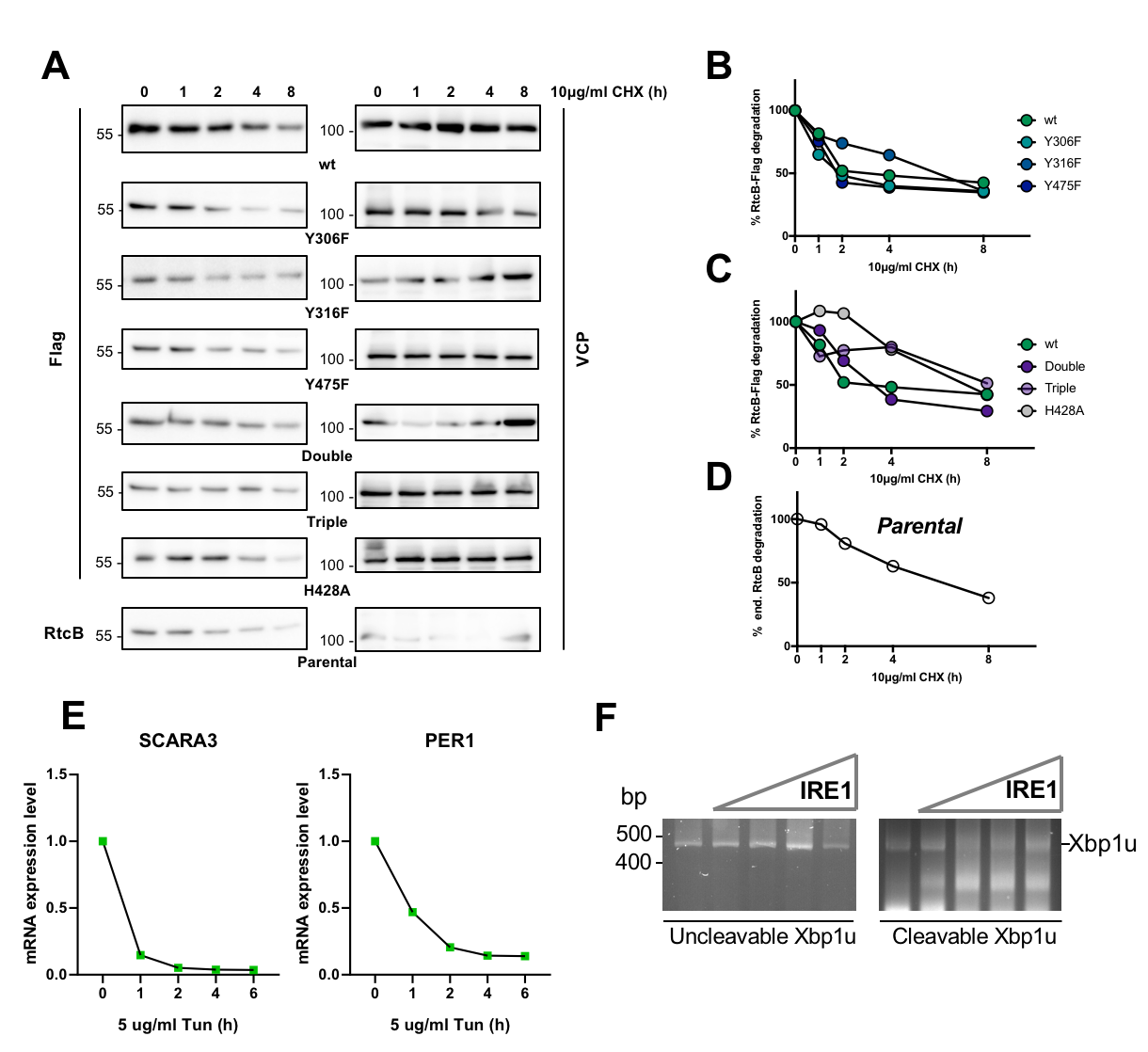

**FIGURE S7**

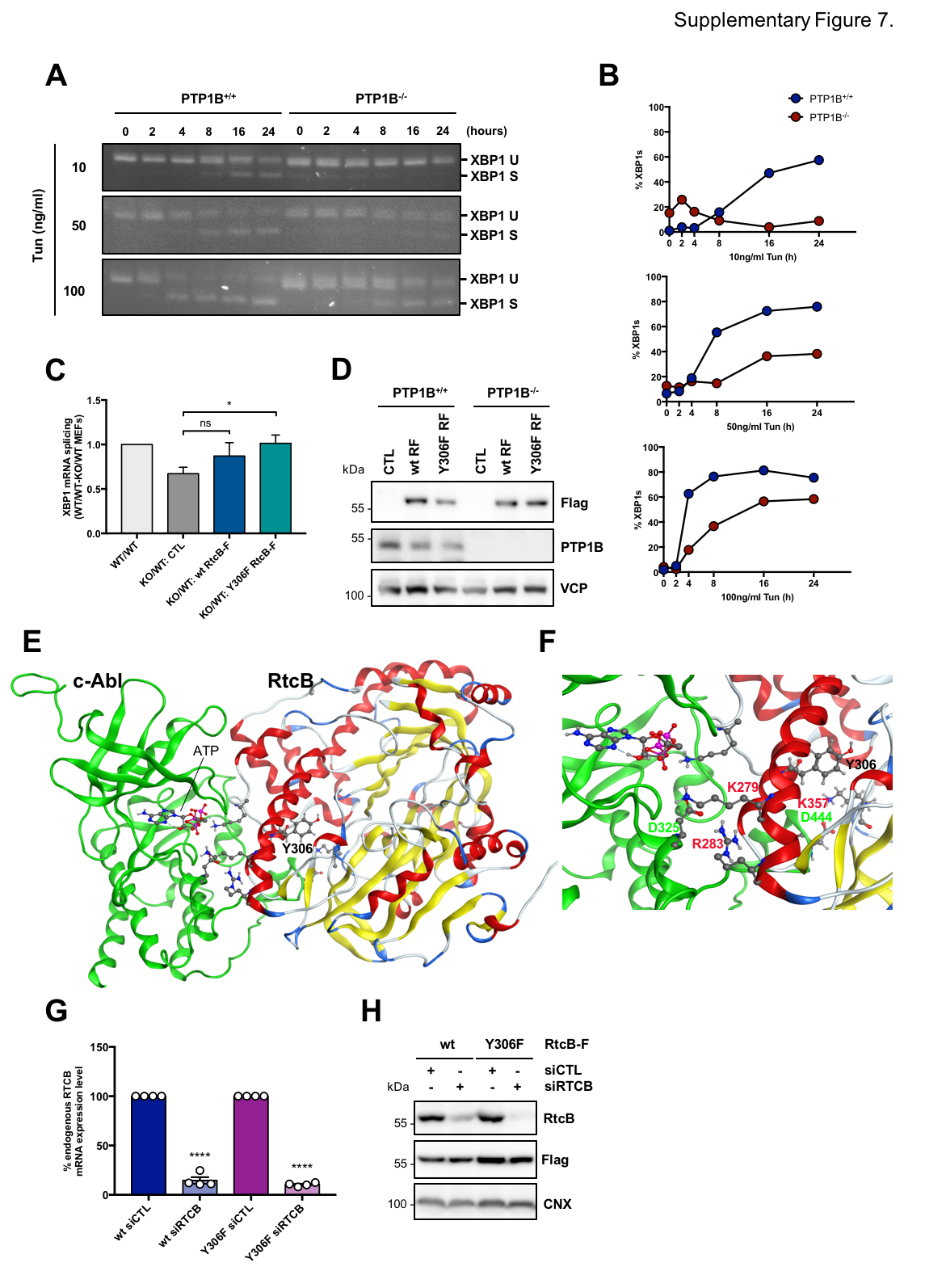

**FIGURE S8**

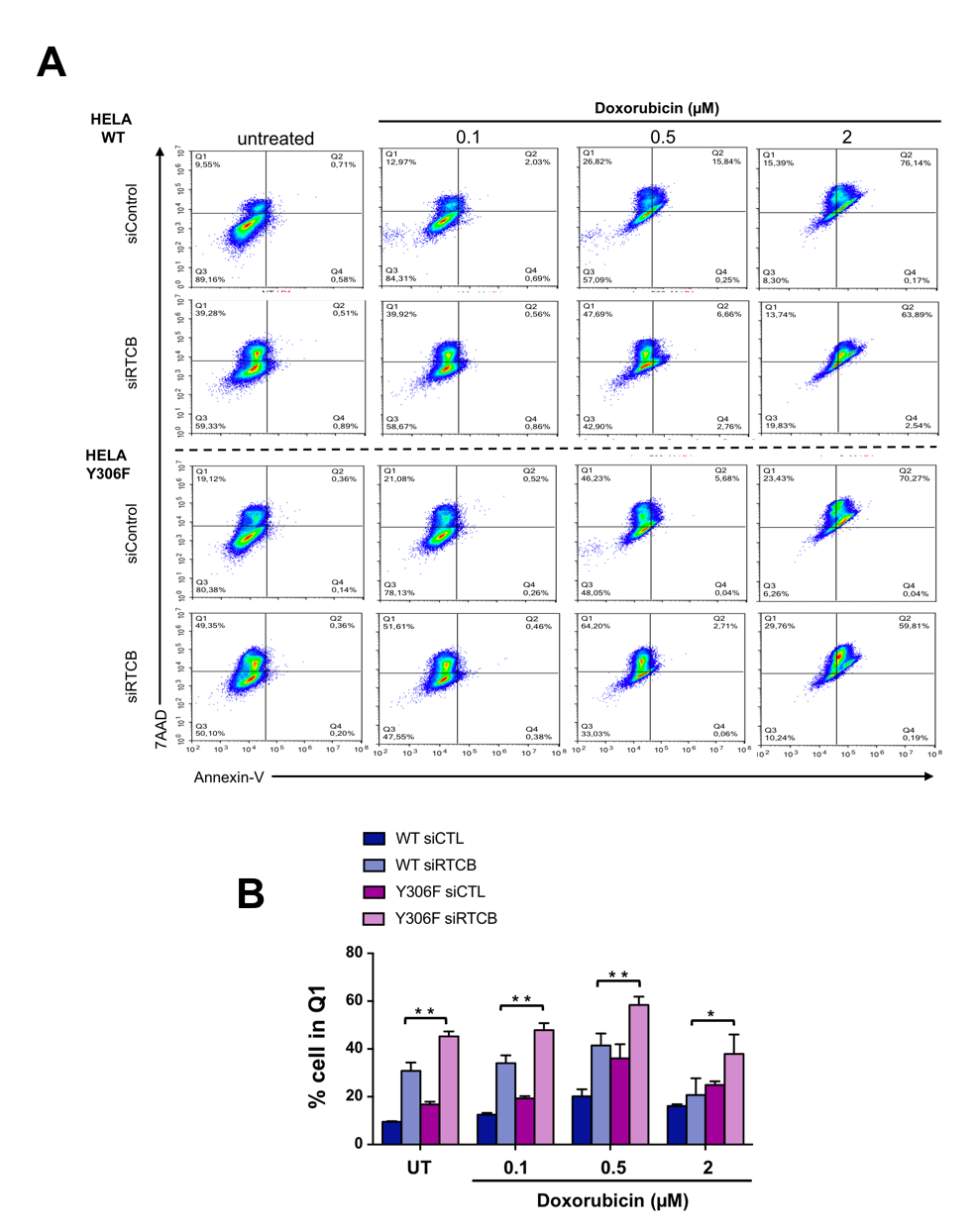

**FIGURE S9**

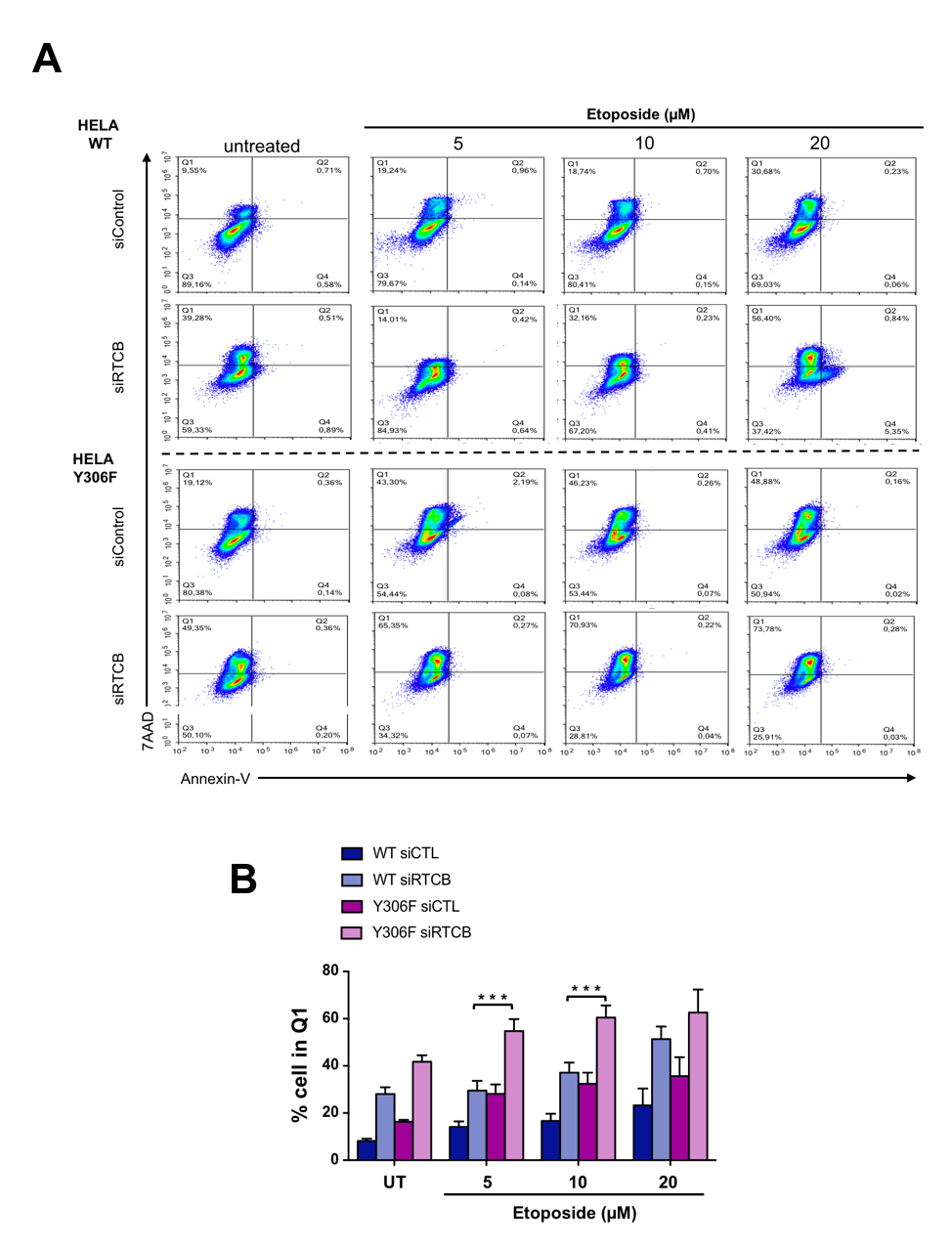

**FIGURE S10**
